# DeepBindGCN: Integrating Molecular Vector Representation with Graph Convolutional Neural Networks for Accurate Protein-Ligand Interaction Prediction

**DOI:** 10.1101/2023.03.16.528593

**Authors:** Haiping Zhang, Konda Mani Saravanan, John Z.H. Zhang

## Abstract

The core of large-scale drug virtual screening is to accurately and efficiently select the binders with high affinity from large libraries of small molecules in which nonbinders are usually dominant. The protein pocket, ligand spatial information, and residue types/atom types play a pivotal role in binding affinity. Here we used the pocket residues or ligand atoms as nodes and constructed edges with the neighboring information to comprehensively represent the protein pocket or ligand information. Moreover, we find that the model with pre-trained molecular vectors performs better than the onehot representation. The main advantage of DeepBindGCN is that it is non-dependent on docking conformation and concisely keeps the spatial information and physical-chemical feature. Notably, the DeepBindGCN_BC has high precision in many DUD.E datasets, and DeepBindGCN_RG achieve a very low RMSE value in most DUD.E datasets. Using TIPE3 and PD-L1 dimer as proof-of-concept examples, we proposed a screening pipeline by integrating DeepBindGCN_BC, DeepBindGCN_RG, and other methods to identify strong binding affinity compounds. In addition, a DeepBindGCN_RG_x model has been used for comparing performance with other methods in PDBbind v.2016 and v.2013 core set. It is the first time that a non-complex dependent model achieves an RMSE value of 1.3843 and Pearson-R value of 0.7719 in the PDBbind v.2016 core set, showing comparable prediction power with the state-of-the-art affinity prediction models that rely upon the 3D complex. Our DeepBindGCN provides a powerful tool to predict the protein-ligand interaction and can be used in many important large-scale virtual screening application scenarios.

## Introduction

Proteins play a key role in most cellular processes, meanwhile ligands can act as mediators of protein and can combat diseases with their physical-chemical properties(Klebe, 2013). However, identifying active compounds experimentally on a large scale is expensive and time-consuming. Hence, the computer aided lead discovery is usually the initial stage of the drug discovery process to reduce the experimental testing burden. Accurately and efficiently predicting the protein-ligand interaction by the computational method is a core component of large-scale drug screening. In recent years, deep learning and machine learning have be widely applied in biology research (Savojardo *et al*., 2018; Z. Chen *et al*., 2021). With the development of deep learning algorithms and increasing protein-ligand interaction data, especially the high resolution atomic structure and experimental binding affinity information, it is possible to apply deep learning to discriminate the binders from nonbinders and predict the affinity. Some affinity prediction models have already been developed, such as pafnucy(Stepniewska-Dziubinska *et al*., 2018), GraphDTA(Nguyen *et al.*, 2021), GAT-Score(Yuan *et al.*, 2021), BAPA(Seo *et al.*, 2021), and AttentionDTA(Zhao *et al*., 2019). Our group also developed DeepBindRG(H. Zhang, Liao, Saravanan, *et al*., 2019) for protein-ligand affinity prediction with the interface atomic contact information as input and DeepBindBC(Zhang, Zhang, *et al*., 2021) for predicting whether protein-ligand complexes are nativelike by creating a large protein-ligand decoy complex set as a negative training set. Moreover, we also developed DFCNN for the preliminary stage of virtual screening since it demonstrates predictable efficiency(H. Zhang, Liao, Cai, *et al*., 2019; Zhang, Lin, *et al*., 2022). Some of our developed models are already applied in drug candidates and target searching, and show huge potential in drug development(Zhang, Li, *et al*., 2021; Zhang *et al*., 2020). However, several limitations still need attention, both in terms of efficiency and accuracy.

The Graph Convolutional Network (GCN) is a kind of deep learning that can use nodes to contain feature information and edges to contain spatial information between nodes, which is a popular method in prediction relationships(S. Zhang *et al*., 2019). GCN is already well applied to predicting the compound property, and molecular fingerprint(Kojima *et al*., 2020; J. Chen *et al*., 2021). Also, the GCN was successfully used for protein-ligand interaction prediction(Nguyen *et al*., 2021; Torng and Altman, 2019). Wen et al. have applied the GCN to predict protein-ligand interactions and achieved encouraging result in the test set. However, they used the DUD-E as a training dataset and only contain 102 receptors, which is very limited diversity in protein information(Torng and Altman, 2019), this strongly suggests their model still has ample improvement space. Its under-trainings on the protein side also can influence its performance significantly. Thin et al. have developed a GCN based protein-ligand prediction model(Nguyen *et al*., 2021), but it used only GCN for the ligand part, and the protein was represented as a sequence, comparing the pocket with spatial information. This sequence lost spatial information and contained much irrelevant information about the protein-ligand binding. Furthermore, Moesser et al. have integrated protein-ligand contact information in ligand-shaped 3D interaction Graphs to improve binding affinity prediction(Moesser *et al*., 2022). Still, it would only be helpful if the protein-ligand complex is available or is accurately predicted by docking.

It should be noted that many deep learning-based protein-ligand affinity prediction models are rarely used in real applications. Even their RMSE value in the testing set seems very small. One major reason is that the affinity model is trained over a binding protein-ligand dataset and doesn’t learn anything about non-binding, while in a real application; the non-binding compounds are dominant during screening over a given target. Hence, purely developing a deep learning-based affinity prediction model is not enough to fulfil the requirement of virtual screening. Developing a model which trained with binding and non-binding data to identify whether protein-ligand was binding is important in the real applications. For instance, we have previous models DFCNN and DeepBindBC to identify whether protein and ligand are binding. These two models successfully helped to identify a given target’s inhibitors with experimental validation in our previous work(Zhang, Zhang, *et al.*, 2022; Zhang, Lin, *et al*., 2022; Zhang, Gong, *et al*., 2022; Zhang *et al*., 2020; Zhang, Li, *et al*., 2021). Moreover, combining the protein-ligand binding prediction model with the affinity prediction model can be more powerful in identifying strong affinity candidates. As aforementioned, hybrid screening has been used to virtualize potential drugs for given targets. However, we still lack a model that can screen over a database size of 100,000~1000,000 accurately and efficiently with the ability to distinguish spatial and physical-chemical features of protein-ligand binding.

In our work, we have used a graph to represent the protein pocket and ligand, respectively, and the GCN model with two inputs and one output to fully train over a large protein-ligand dataset PDBbind. The diversified structure database PDBbind guarantees the robustness of model performance. We also evaluate the model performance using the known binding and nonbinding data. We also show its application in drug candidate screening for target TIPE3 and PD-L1 dimers. Our result shows DeepBindGCN can be a valuable tool to rapidly identify reliable, strong binding protein-ligand pairs and can be an essential component of a hybrid large scale screening pipeline.

## Method

### Data preparation

The training data is downloaded from PDBbind2019. The protein pocket was defined as a cutoff value within the known ligand (any atom in the residue within the cutoff value of the ligand will keep the residue as pocket residue). We tested cutoff values of 0.6 nm and 0.8nm in this work. The ligands were represented as molecule graphs by converting the SMILES code to its corresponding molecular graph and extracting atomic features using the open-source chemical informatics software RDKit(Landrum, 2006).

The pocket was represented as a graph by defining the residues as nodes and contacting residue pairs as edges (the cutoff was set as 0.5 nm). We have tested onehot and molecular vector representations for the node residue, respectively. A pretrained mol2vec model generated the molecular vector.

### The dataset for a binary classification task

Through cross-combination, we obtain 52200 protein-ligand pairs as a negative dataset and divide them into 45000 as training negative data and 7200 as testing negative data. From the PDBbind2019 dataset, we obtained a total of 17400 proteinligand as positive data, divided into 15000 as training positive data, and 2400 as testing data. During the training, the positive training and testing data are used 3 times to keep the positive and negative data balanced.

### The dataset for the affinity prediction task

We obtained 16956 protein-ligand datasets with affinity from PDBbind2019 and divided them into 15000 training and 1956 test datasets. In the PDBbind v2019 dataset, the binding affinities of protein-ligand complexes were provided with Ki, Kd, and IC50. We transformed the binding affinities into pKa using the following equation:

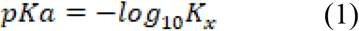

where *K_x_* represents IC50, *K_i_*, or *K_d_*.

### Pre-train 30-dimension molecular vector to represent residues in pocket

We downloaded 9,216,175 onstock compounds from the ZINC15 database as a training dataset, the mol2vec was used to do the training, and we finally obtained a model that can generate a vector for each given chemical group, here we set the vector dimension to 30. The obtained model was used to generate the vector of the 20 residues by adding the chemical group vectors within each residue.

### Model construction

The model structure is shown in Figure 1. It has two inputs (drug–target pair) and one output structure. The ligand and pocket graphic information flow into the two layers of the graphic network. Then, the output of two graphic networks is merged into fully connected layers. The final output was one node. The binary prediction uses the sigmoid activation function, which gives a value range of 0~1; for the affinity prediction, the output uses linear activation, which is a continuous measurement of binding affinity for that pair.

**Figure 1.**
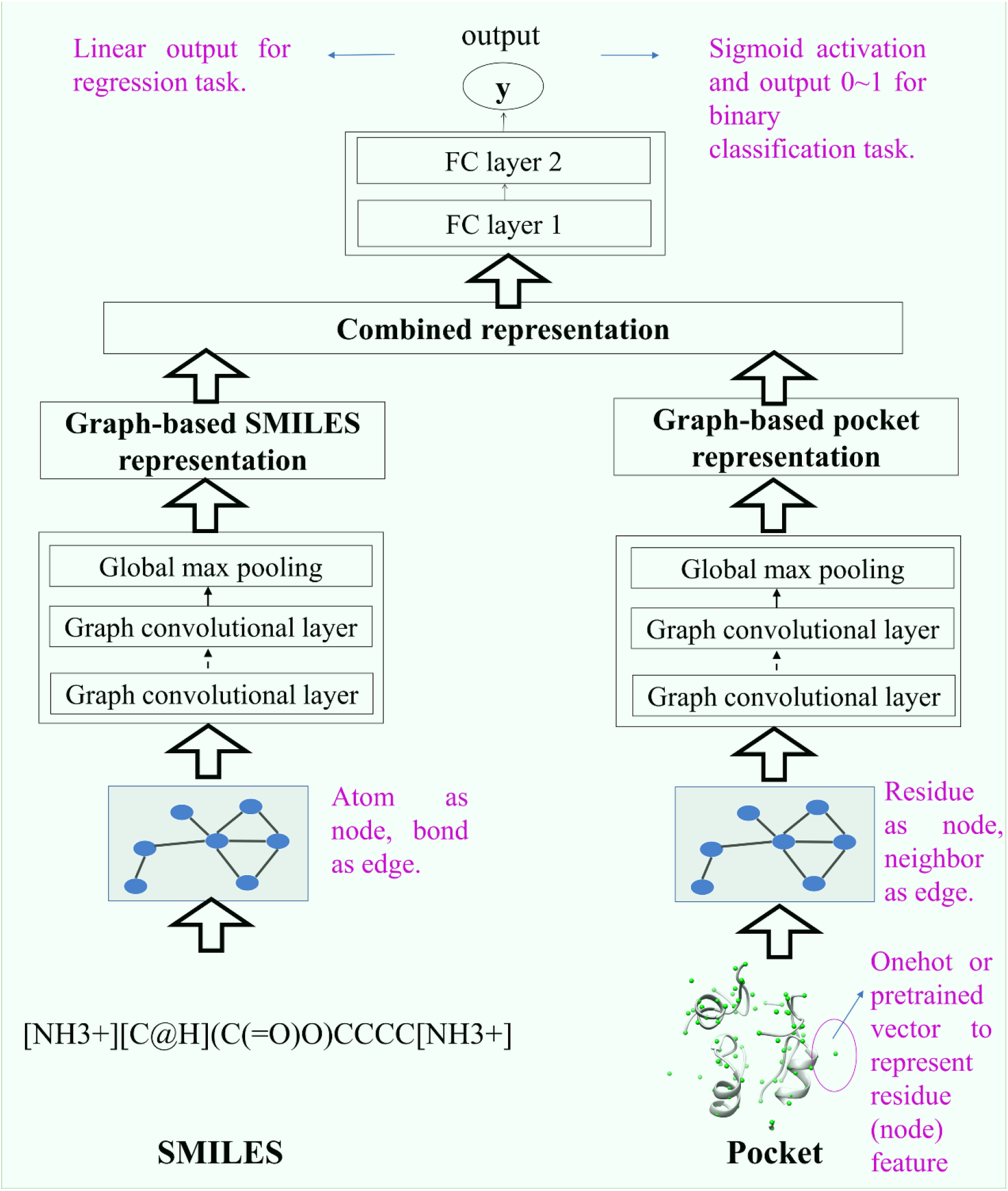
The architecture of the DeepBindGCN model.

### Model training

The torch_geometric module was used to create input data and construct the graphic neural network. The input data was saved in PyTorch, InMemoryDataset format. The PyTorch was used to do the training. The number of epochs that we finally chose was based on the performance convergences on the test set.

### Model performance compared with other methods on the DUD.E dataset

We have downloaded 102 therapeutic-related proteins and their corresponding active and inactive compounds from the DUD.E dataset(Mysinger *et al*., 2012). Those data were processed into the input format and used as extra testing set to examine our model performance. The performance matrix AUC, MCC, Accuracy, Precision, and TPR were used to validate the BC model, and the rmse, mse, pearson correlation, spearman correlation, and Concordance Index (CI) were used to validate the RG model.

### Virtual screening of candidates against two targets (TIPE3 and PD-L1 dimer)

The atomic coordinates of TIPE3 were retrieved from PDB with id 4Q9V(Fayngerts *et al*., 2014). The TIPE3-ligand complex was modeled by the cofactor method in https://zhanggroup.org/COFACTOR/ web server(Roy *et al*., 2012). The PD-L1 dimer was retrieved from PDB with id 5N2D(Guzik *et al*., 2017), these PDB structures already contain ligands. The pocket was extracted as 0.8 nm from the predicted or known ligands. The dataset Chemdiv with the size of 1,507,824 compounds, was used as a virtual screening dataset.

### Tools used in the analysis

The USCF Chimera, VMD, Schrödinger, pymol, and Discovery Studio Visualizer 2019 were used to generate the structure and to visualize the 2D protein-ligand interactions(Pettersen *et al*., 2004; Humphrey *et al*., 1996; Visualizer, 2005). Clusfps (https://github.com/kaiwang0112006/clusfps), which depends on RDKit(Landrum, 2006), was used to cluster the drugs in the dataset. The drug fingerprint was used as an input, with the algorithm of Murtagh(Murtagh and Contreras, 2012) being used for clustering candidates into 6 groups.

## Results

The DeepBindGCN_BC and DeepBindGCN_RG workflow is shown in Figure S1, we observed that during the application, their input preparation, and model architecture are highly consistent, except that one is output 0~1 for binary classification, and the other is output continuous value for affinity prediction.

### The performance of DeepBindGCN_BC and DeepBindGCN_RG on training and test set

The AUC, TPR, Precision, and accuracy of the training set and test set over the 2000 epoch training for the DeepBindGCN_BC are recorded and shown in Figure S2 and Table S1. The AUC values fall around 0.86~0.87 and 0.84~0.85 after 400 epochs when using pocket cutoff value 0.6nm and 0.8 nm, respectively, indicating the training has fully converged in epoch 2000. The result also shows that the DeepBindGCN_BC performs better on the testing set when using a pocket cutoff of 0.8nm according to the performance metrics AUC, TPR, precision, and accuracy. For instance, the DeepBindGCN_BC has AUC, TPR, precision, and accuracy values of DeepBindGCN_BC with cutoff 0.6nm at epoch 2000 are 0.8788, 0.6863, 0.6767, and 0.8396, respectively, corresponding to values 0.8537, 0.6175, 0.6552, and 0.8231 when with pocket cutoff 0.8nm, which all demonstrate slight better performance.

The rmse, mse, pearson correlation, spearman correlation, and Concordance Index (CI, the larger, the better) of the training set and test set over the 2000 epoch training for the DeepBindGCN_RG are shown in Figure S3, Table S2. We noted that the RMSE has stayed around values 1.3 and 1.1~1.3 after 400 epochs when using pocket cutoff values 0.6nm and 0.8 nm respectively, indicating that the training has fully converged. The DeepBindGCN_RG has better performance with a pocket cutoff of 0.8nm compared to a pocket cutoff of 0.6nm according to the performance metrics rmse, mse, pearson correlation, spearman correlation, and CI. For instance, DeepBindGCN_RG with the pocket cutoff of 0.8nm has rmse, mse, pearson correlation, spearman correlation, and CI values of 1.2107, 1.4657, 0.7518, 0.7410, and 0.7756 in epoch 2000, respectively, corresponding to values of 1.3361, 1.7852, 0.7141, 0.7098, and 0.7628 when the pocket cutoff is 0.6nm, which all demonstrate slight better performance in pocket cutoff 0.8nm.

Interestingly, we found that the pocket cutoff value of 0.6 nm has a better performance for the DeepBindGCN_BC, while the cutoff value of 0.8 nm has a better performance for the DeepBindGCN_RG. This suggests that the close contact ligand and residue information is enough to accurately predict whether protein-ligand is binding, and long-range contact information sometimes may mislead its prediction. However, long-range pocket residue information is also important to accurately predict how strong protein-ligand is binding. To accurately estimate the binding affinity, most of the residues that have contributed to the binding should be considered. Notably, we apply a pocket cutoff of 0.6 nm for DeepBindGCN_BC and apply a pocket cutoff of 0.8 nm for the DeepBindGCN_RG in the rest of the work.

### The performance of DeepBindGCN_BC and DeepBindGCN_RG on the DUD.E dataset

We have considered experimental known inactive and active protein-compound pairs or protein-compounds affinity information from the DUD.E dataset for our model extra testing set. Precision is widely acknowledged to be an important performance metric in large-scale virtual screening applications. The performances of DeepBindGCN_BC and DeepBindGCN_RG on some DUD.E datasets are listed in Table S3 and Table S4, respectively. We noticed that DeepBindGCN has a very high precision (>0.9) over more than half of the cases from the DUD.E datasets, as shown in Table 1. It should also be noted that many other performance metrics are not good for DeepBindGCN in many cases, as shown in Tables 1 and S3. Some protein-ligand datasets are predicted into all 0 values, which indicate no binding. A possible explanation is that the binding pocket we selected cannot guarantee exactly binding with those ligands. Also, the data may contain some false positive experimental results since there are no crystal structures as strong proof of binding. To sum up, the high precision of DeepBindGCN_BC in DUD.E data guarantees that the selected compounds from large-scale virtual screening are likely to be binders.

**Table 1.**
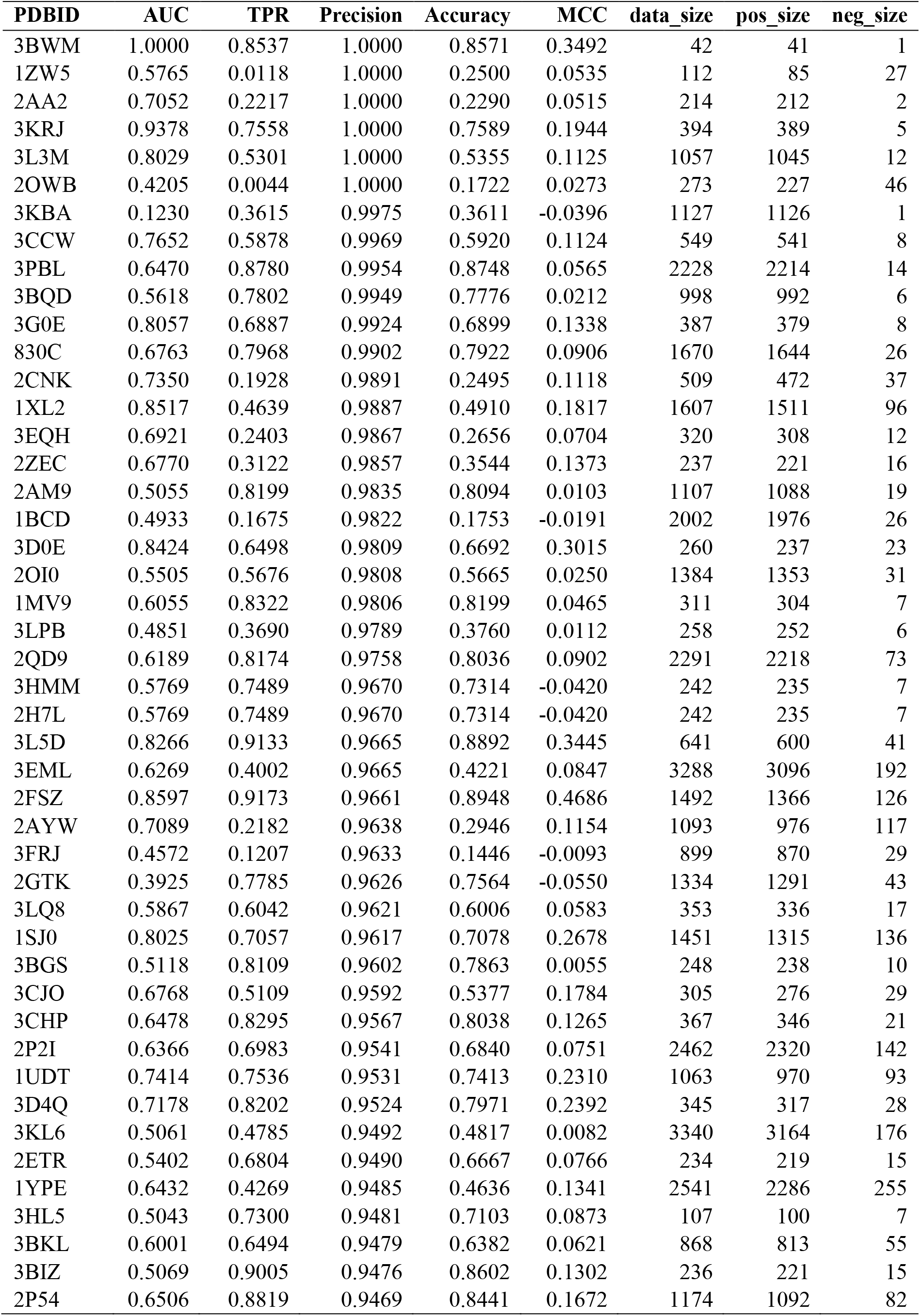
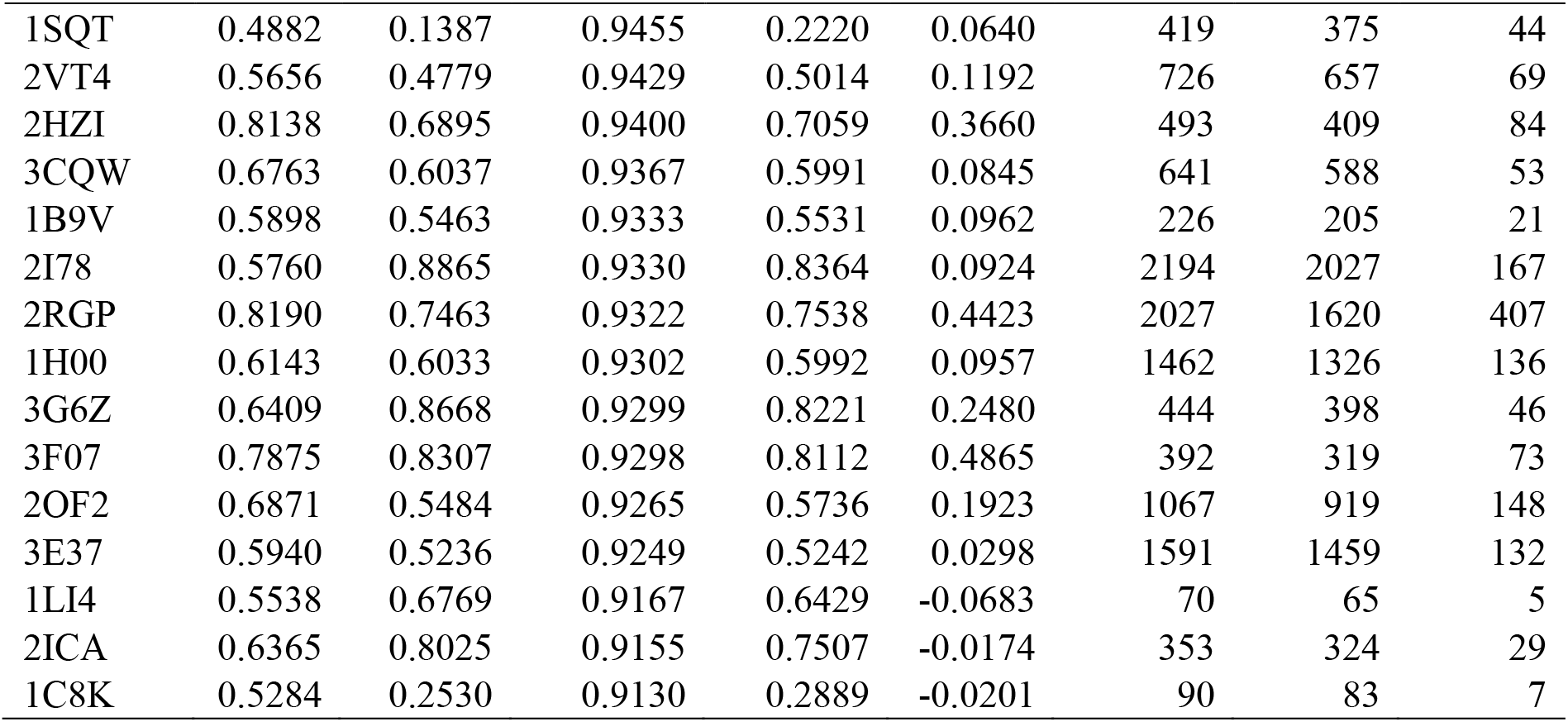
The performance of DeepBindGCN_BC on some of the DUD.E datasets with precision values larger than 0.9.

We also tested the DeepBindGCN_RG on the DUD.E dataset with affinity values, shown in Table S4. Interestingly, DeepBindGCN_RG performs well over most datasets in terms of rmse values. The average rmse of 102 therapeutic targets related datasets has reached 1.1893. We can see that more than 65 protein target-related dataset has rmse smaller than 1.2, as shown in Table 2, which is extremely accurate compared to most of the current affinity prediction methods. On the other hand, the pearson correlation, spearman correlation, and CI also demonstrate that prediction and experimental measurement usually have a weak correlation. We believe this is mainly because for each dataset, many compounds with affinity have similar structures, hence making the model extremely challenging to detect the slightly binding affinity difference. The low rmse and mse can guarantee that the DeepBindGCN_RG can correctly select strong affinity binders out of the abundant candidates from DeepBindGCN_BC.

**Table 2.**
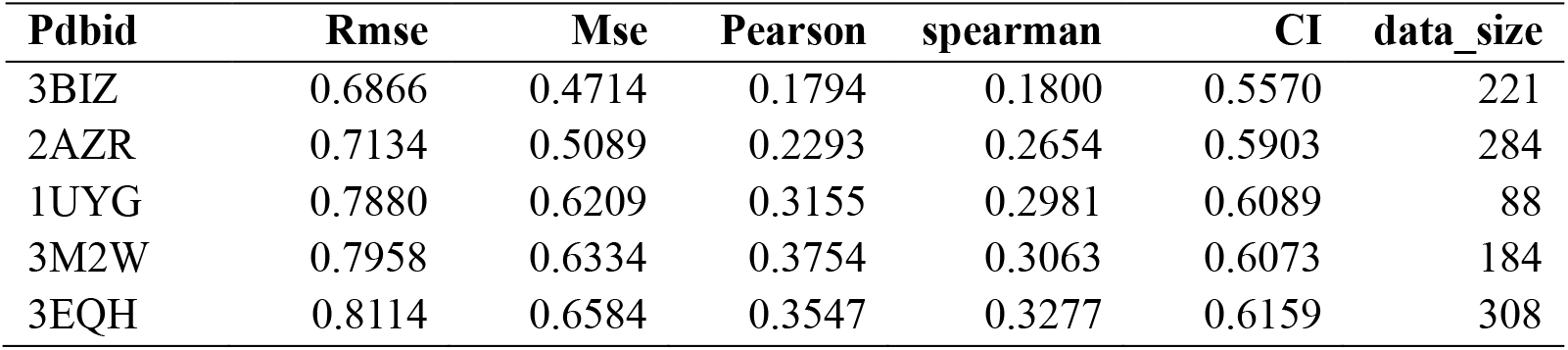
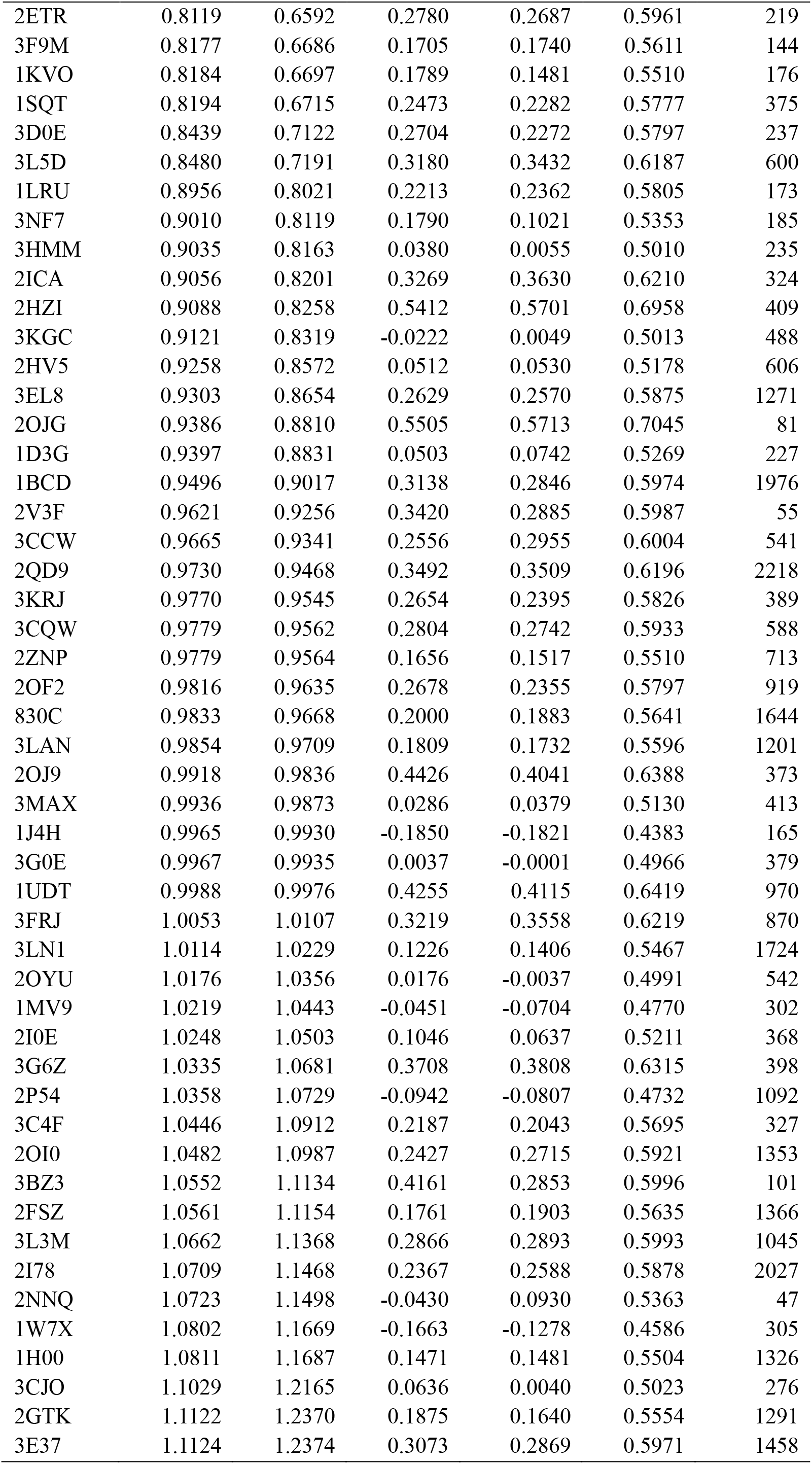
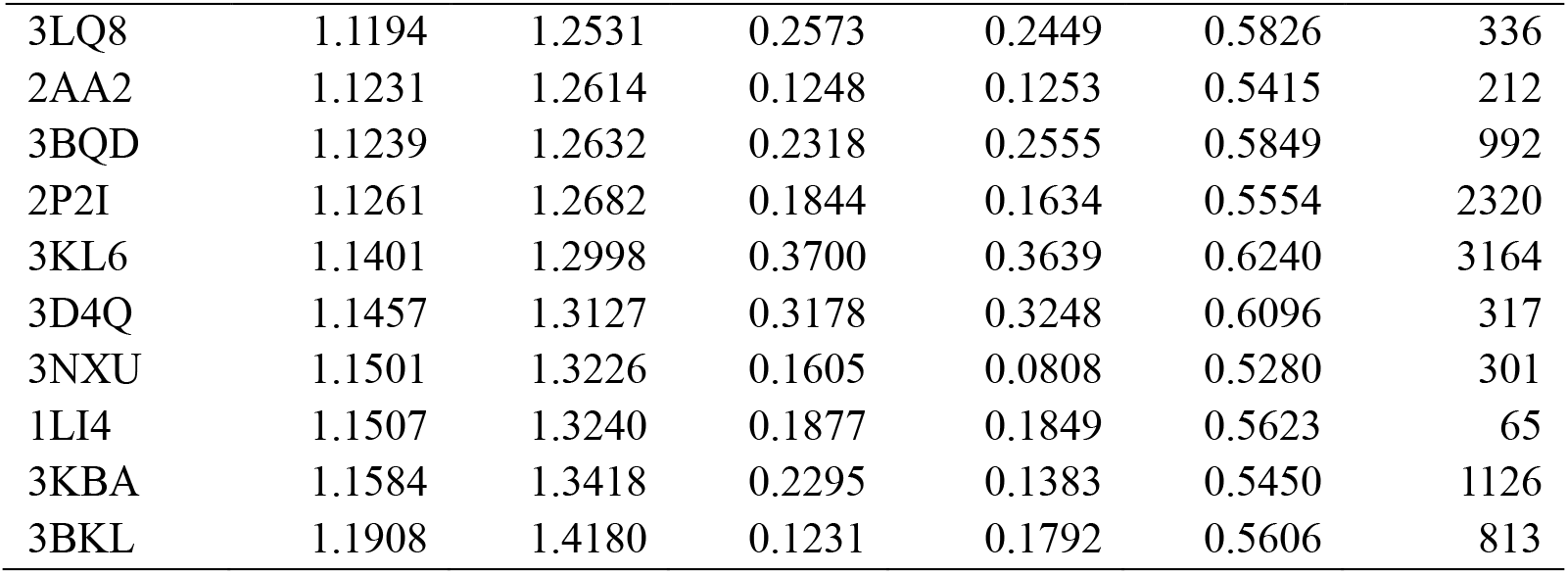
The performance of DeepBindGCN_RG on some DUD.E datasets with rmse smaller than 1.2.

### Virtual screening by DeepBindGCN against TIPE3 and PD-L1 dimer as self-concept-approve examples

The screening application diagram with screening against TIPE3 as an example is illustrated in Figure 2, which integrates many different methods, including DeepBindGCN, docking, MD simulation, and Metadynamics-based binding free energy landscape calculation. The MD and Metadynamics simulation details are described in Supplementary material section 2.

**Figure 2.**
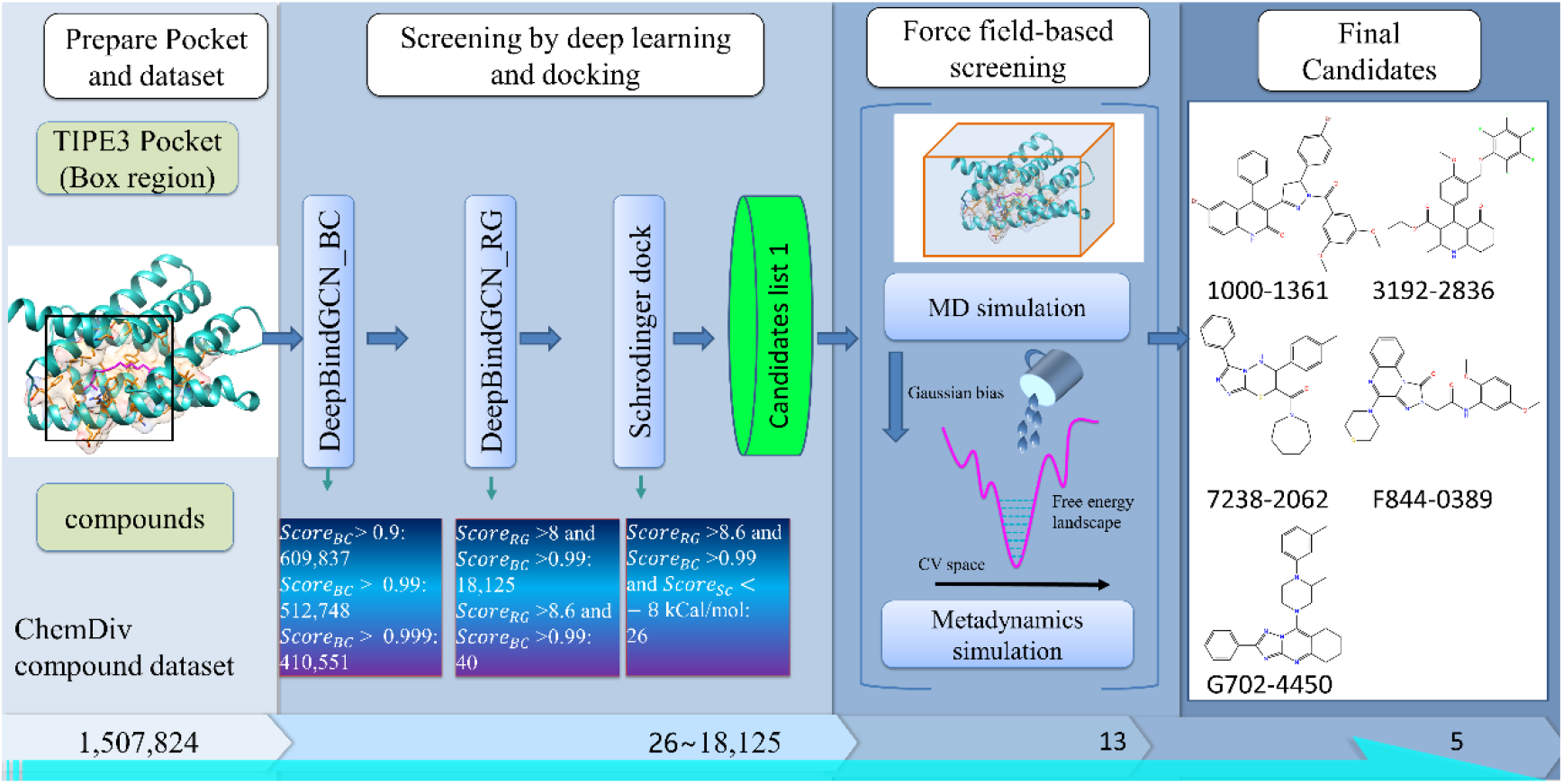
The virtual screening procedure integrates DeepBindGCN models with other methods to identify highly reliable drug candidates for TIPE3.

The TIPE3 is a transfer protein for lipid second messengers and is upregulated in human lung cancer tissues (Fayngerts *et al*., 2014). Recent research reveals its important role in cancer proliferation, which is believed to be a novel cancer therapeutic target(Li *et al*., 2021). However, there are still no effective compounds that can inhibit its function. In this work, we obtain 40 compound candidates with DeepBindGCN_BC score > 0.99 and DeepBindGCN_RG > 9, shown in Table 3. We also docked these candidates with TIPE3 by Schrödinger software to obtain the potential binding conformations. The docking score is listed in Table 3.

**Table 3.**
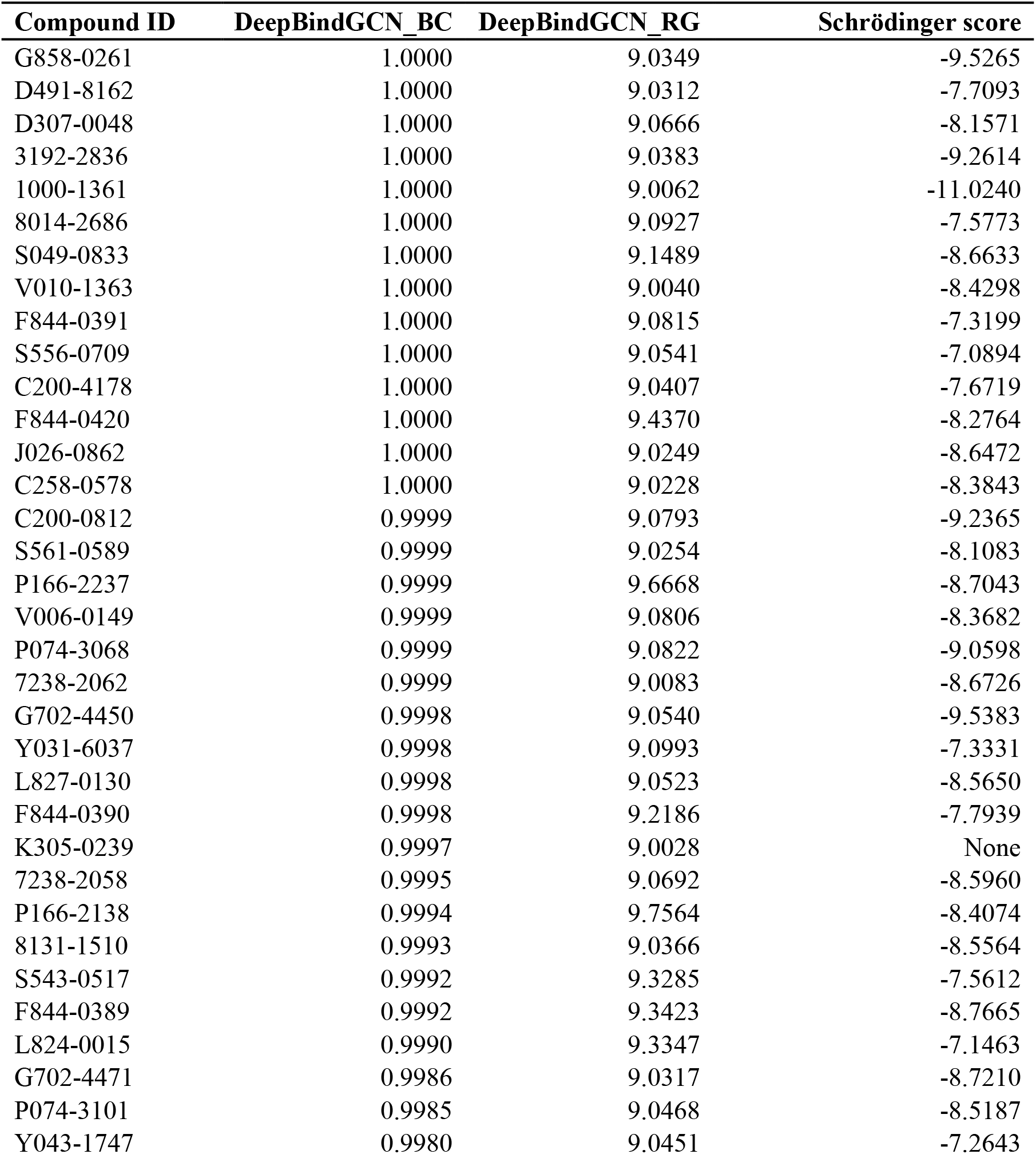
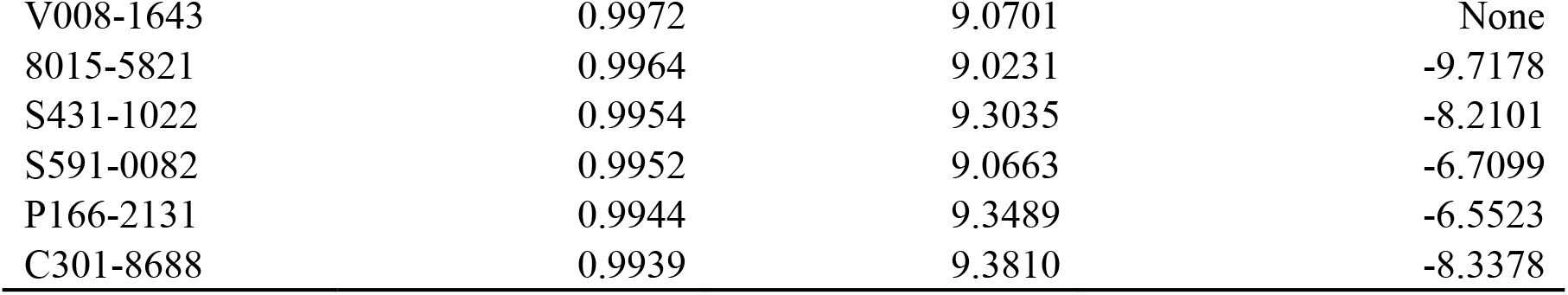
The top predicted candidates from DeepBindGCN_BC and DeepBindGCN_RG for the TIPE3.

| Compound ID | DeepBindGCN_BC | DeepBindGCN_RG | Schrödinger score |
| --- | --- | --- | --- |
| G858-0261 | 1.0000 | 9.0349 | -9.5265 |
| D491-8162 | 1.0000 | 9.0312 | -7.7093 |
| D307-0048 | 1.0000 | 9.0666 | -8.1571 |
| 3192-2836 | 1.0000 | 9.0383 | -9.2614 |
| 1000-1361 | 1.0000 | 9.0062 | -11.0240 |
| 8014-2686 | 1.0000 | 9.0927 | -7.5773 |
| S049-0833 | 1.0000 | 9.1489 | -8.6633 |
| V010-1363 | 1.0000 | 9.0040 | -8.4298 |
| F844-0391 | 1.0000 | 9.0815 | -7.3199 |
| S556-0709 | 1.0000 | 9.0541 | -7.0894 |
| C200-4178 | 1.0000 | 9.0407 | -7.6719 |
| F844-0420 | 1.0000 | 9.4370 | -8.2764 |
| J026-0862 | 1.0000 | 9.0249 | -8.6472 |
| C258-0578 | 1.0000 | 9.0228 | -8.3843 |
| C200-0812 | 0.9999 | 9.0793 | -9.2365 |
| S561-0589 | 0.9999 | 9.0254 | -8.1083 |
| P166-2237 | 0.9999 | 9.6668 | -8.7043 |
| V006-0149 | 0.9999 | 9.0806 | -8.3682 |
| P074-3068 | 0.9999 | 9.0822 | -9.0598 |
| 7238-2062 | 0.9999 | 9.0083 | -8.6726 |
| G702-4450 | 0.9998 | 9.0540 | -9.5383 |
| Y031-6037 | 0.9998 | 9.0993 | -7.3331 |
| L827-0130 | 0.9998 | 9.0523 | -8.5650 |
| F844-0390 | 0.9998 | 9.2186 | -7.7939 |
| K305-0239 | 0.9997 | 9.0028 | None |
| 7238-2058 | 0.9995 | 9.0692 | -8.5960 |
| P166-2138 | 0.9994 | 9.7564 | -8.4074 |
| 8131-1510 | 0.9993 | 9.0366 | -8.5564 |
| S543-0517 | 0.9992 | 9.3285 | -7.5612 |
| F844-0389 | 0.9992 | 9.3423 | -8.7665 |
| L824-0015 | 0.9990 | 9.3347 | -7.1463 |
| G702-4471 | 0.9986 | 9.0317 | -8.7210 |
| P074-3101 | 0.9985 | 9.0468 | -8.5187 |
| Y043-1747 | 0.9980 | 9.0451 | -7.2643 |
| V008-1643 | 0.9972 | 9.0701 | None |
| 8015-5821 | 0.9964 | 9.0231 | -9.7178 |
| S431-1022 | 0.9954 | 9.3035 | -8.2101 |
| S591-0082 | 0.9952 | 9.0663 | -6.7099 |
| P166-2131 | 0.9944 | 9.3489 | -6.5523 |
| C301-8688 | 0.9939 | 9.3810 | -8.3378 |

For the convenience of analysis, we have clustered the candidate’s list into six groups, and the structures of cluster center compounds are shown in Figure S4. We observed clusters 1 and 2 have the largest number of group members. Notably, the cluster center structure contains several benzene-like substructures, indicating that pi-related interactions may be necessary for strong binding with TIPE3. We also notice that the representative structures of clusters 1, 2, 3, 4, and 5 have a linear shape, indicating that the linear shape molecules may easier enter the binding cavity and achieve tightly binding. Also, all the representative structures are relatively flat, which may help enter the binding cavity more easily.

To further explore the predicted TIPE3’s interaction details with the representative structures, we have plotted its 3D and 2D pocket-ligand interaction details, shown in Figure 3. Consistent with our previous assumption, we observed that most interactions are strongly maintained by Pi-related interaction. Only F844-0389 has formed one hydrogen bond with TIPE3, while there are many pi-related interactions for most of these compounds with TIPE3, indicating the hydrogen bond is may not the dominant force for tightly TIPE3 binding. Compound-induced dimerization of PD-L1 is an effective way to prevent PD-L1-PD-1 binding, leading to inhibiting cancer cell proliferation. We have carried out DeepBindGCN screening over the compounds database. The compounds with DeepBindGCN_BC > 0.99 and DeepBindGCN_RG >8.6 were selected as candidates, shown in Table S5.

**Figure 3.**
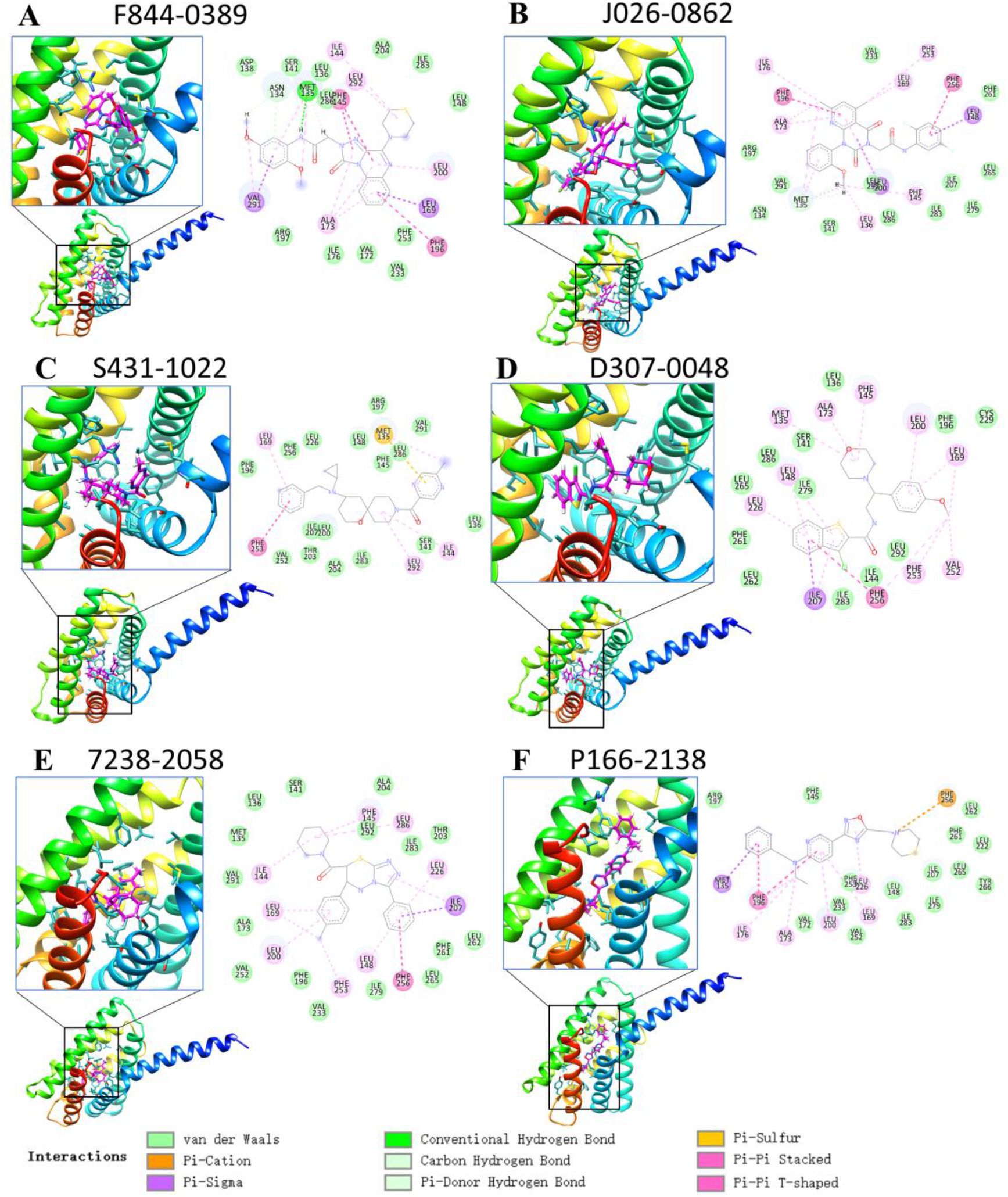
The snapshot and 2D plot of TIPE3 with representative cluster center compounds from docking.

We obtain 6 representative structures by clustering the candidates into six groups, as shown in Figure S5. Cluster 5 has the largest group members. The representative structures of clusters 1, 2 and 3 have a similar shape, while the representative clusters 3, 4, and 5 share similar linear shapes. Interestingly, except representative structure 2, all other 5 representative structures are compounds with the pentacyclic ring. The 2D interaction of the predicted representative compounds with PD-L1 dimmer from Schrödinger docking is shown in Figure S6. Most compounds interact with the PD-L1 pocket, including hydrogen bonds, Pi-related interaction, salt bridge interaction, etc. It should be noted that Schrödinger has not successfully docked K305-0238 and E955-0720 into the selected PD-L1 pocket.

We further carried out MD and Metadynamics simulations to check the binding stability of the predicted protein-ligand pairs. The candidates that show favorable binding with the 3 targets according to the free energy landscape from the metadynamics simulation are selected to further analysis, as shown in Figure S7.

We noticed that except F844-0389 (RMSD around 0.3~0.5), the calculated RMSD of these selected candidates for the TIPE3 have very small values (around 0.1~0.3nm) and low fluctuation, as shown in Figure S8, indicating the candidates have very stable binding. The protein-compound interaction details of the last frame from the MD simulation are shown in Figure 4.

**Figure 4.**
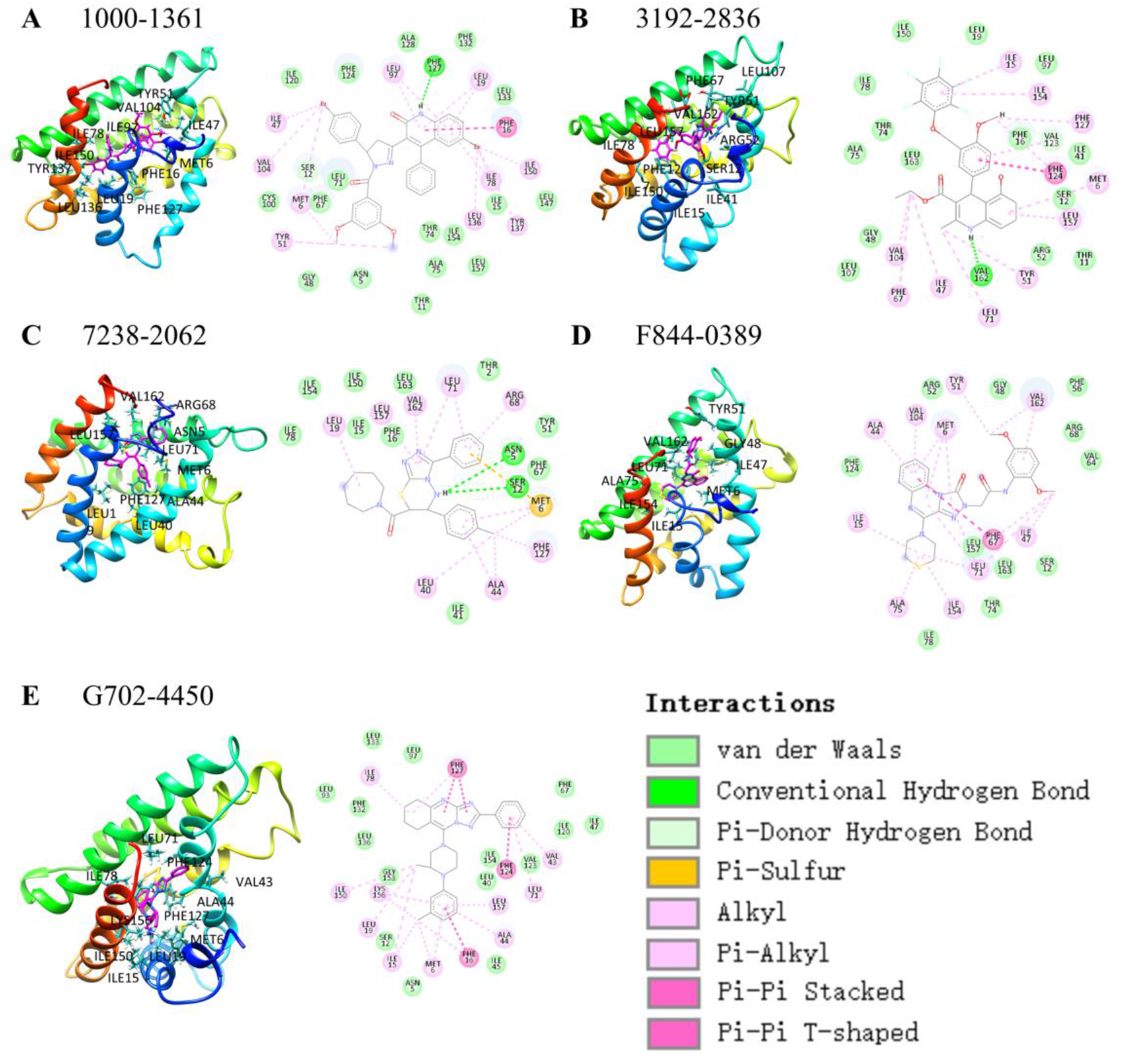
The TIPE3 interaction details with candidate compounds for the last frame from the MD simulation.

The RMSD of the selected candidates for the PD-L1 dimer is shown in Figure S9. Notably, the RMSD values of 4376-0091 and P392-2143 have very small values (around 0.1~0.2nm), indicating their binding is stable. The interaction details of selected candidates with PD-L1 dimer are shown in Figure 5. We observed that the binding pocket contains many non-polar residues, and the interaction between PD-L1 dimer with compounds is dominant with hydrophobic interactions, therefore, this confirms that the compounds act as a molecular glue to promote and stable the PD-L1 dimerization.

**Figure 5.**
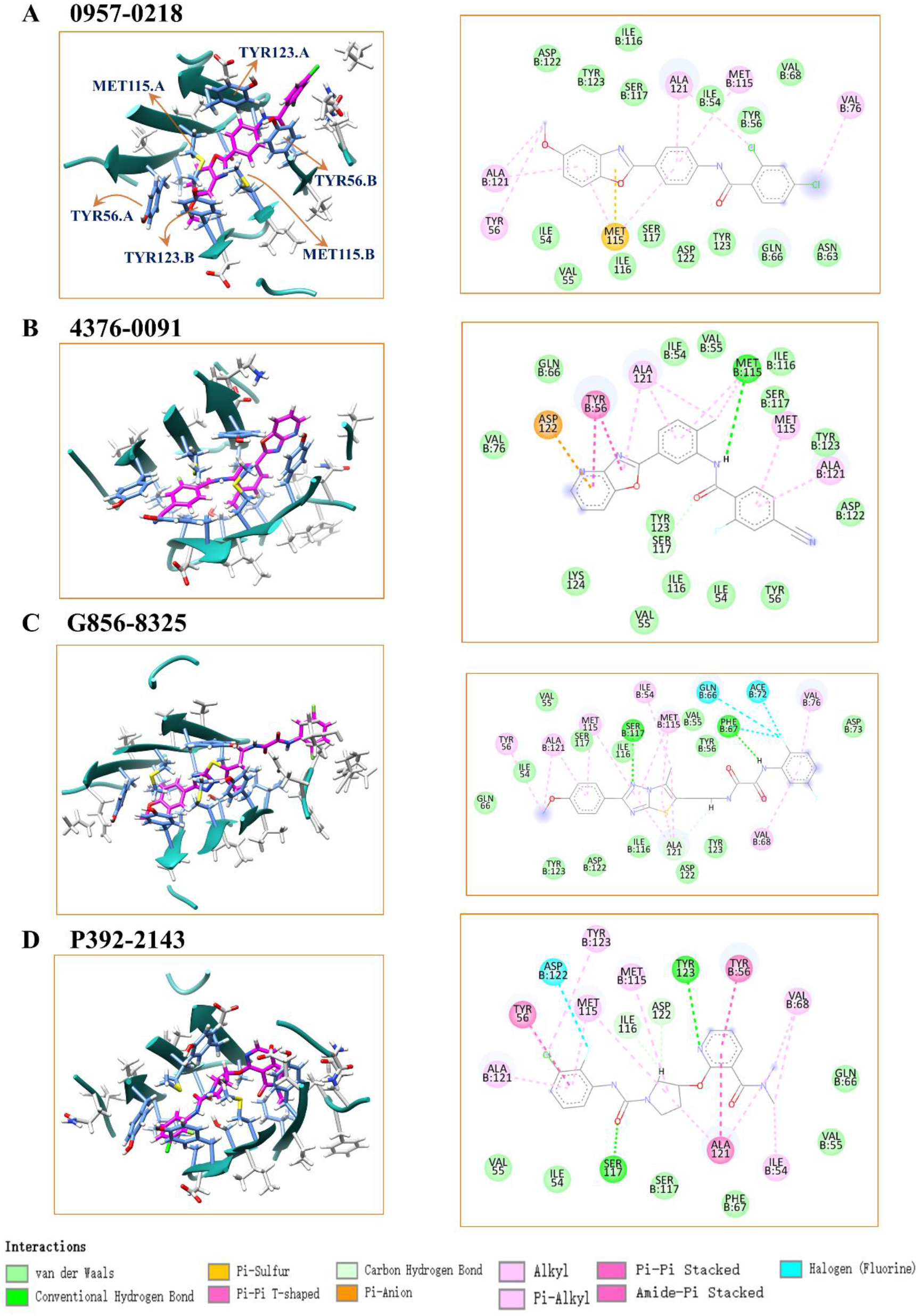
The PD-L1 dimer interaction details with candidate compounds for the last frame from the MD simulation.

## Discussion

The proposed GCN-based model is extremely efficient compared to traditional docking and deep learning-based methods. Since it does not depend on the proteinligand complex, it can save time and resources to preprocess the input by docking. In many other complex structure-based models, most of the time is spent for exploring binding conformation, and the prediction would be highly unreliable if the binding conformations are incorrect. By using the pre-trained molecular vector to represent the residues, the GCN-based model has an obvious improvement, indicating our model can identify physical-chemical features and spatial information. The model’s performance is good on the DUD.E dataset, indicating it’s highly advantageous in real applications. This model has great potential as a core component of large-scale virtual screening. The method is strongly complementary to many existing methods, such as docking, MD simulation, and other deep learning methods; hence can easily be integrated into a hybrid screening strategy. The methods can also be used to screen de novo compounds by combining them with molecular generative models, similar to our previous work(Zhang, Saravanan, *et al*., 2022).

To check its efficiency in virtual screening, we tested its time spent in virtual screening. With CUDA acceleration, we find DeepBindGCN_BC and DeepBindGCN_RG spent about 45.5s and 22.2s to complete the prediction of 50000 protein-ligand pairs, respectively, with an Intel CPU cores (2.00 GHz) and a GeForce RTX 2080 Ti GPU card. With only CPU, it takes about 57.8s and 61.9s for DeepBindGCN_BC and DeepBindGCN_RG to finish the prediction of 50000 proteinligand pairs, respectively, with 40 Intel CPU core (2.00 GHz). This indicates that DeepBindGCN_BC or DeepBindGCN_RG only need 0.0004~0.0012s to complete a prediction, which is at least ten thousand times faster than traditional docking (which usually takes tens of seconds to several minutes) or docking-dependent deep learningbased protein-ligand affinity prediction method. In summary, large-scale virtual screening would greatly benefit from DeepBindGCN’s efficiency.

To compare the performance of the DeepBindGCN_RG-like model with other affinity prediction models on the PDBBIND core set, we have trained a DeepBindGCN_RG_x model over datasets without PDBBIND core set2013 and 2016 (CASF-2016). The training details are in supplementary material section 1. The performance of DeepBindGCN_RG_x with different epochs on PDBBIND core sets 2013 and 2016 (CASF-2016) are shown in Tables S6 and S7. We can see the model has the best performance with epoch 1700 for both datasets. Hence, we are using a model with a 1700 epoch as the final model. Since many other protein-ligand affinity prediction models have widely tested these two datasets, we collected other methods’ performance from literature reports and showed them in Table 4. Those methods used for comparison include KDEEP(Jiménez *et al*., 2018), Pafnucy(Stepniewska-Dziubinska *et al*., 2018), midlevel fusion(Jones *et al*., 2021), GraphBAR(Son and Kim, 2021), AK-score-ensemble(Kwon *et al.*, 2020), DeepAtom(Li *et al.*, 2019), PointNet(B)(Wang *et al.*, 2022), PointTransform(B)(Wang *et al.*, 2022), AEScore(Meli *et al*., 2021), ResAtom-Score(Y. Wang *et al*., 2021), DEELIG(Ahmed *et al*., 2021), PIGNet (ensemble)(Moon *et al*., 2022), BAPA(Seo *et al*., 2021), SE-OnionNet(S. Wang *et al*., 2021), DeepBindRG(H. Zhang, Liao, Saravanan, *et al*.,2019). We can see that our DeepBindGCN_RG_x has comparable performance with most state-of-art models. We noted that some methods have better RMSE or R-value than our DeepBindGCN_RG_x, but they all have utilized interface information of crystal 3D structure of the protein-ligand complex. Moreover, only our method in Table 4 is independent of the protein-ligand complex, while others depend on the experimental complex. The experimental complex is unavailable in a real application, and the protein-ligand complex is obtained by docking. The method will perform poorly in such a scenario due to some unreliable docking conformation. However, our method’s performance is independent of the protein-ligand complex, and its performance would be stable in such a real application. Its good performance in the DUD.E dataset also strongly supports this assumption. It is the first time that a deep learning-based model has achieved a rmse value of 1.3322 and Pearson R-value of 0.7922 in PDBbind v.2016 core set without any 3D protein-ligand complex. This affinity prediction model is valuable in a wide range of real-case virtual screening applications. In contrast, most current affinity prediction models are rarely used in real applications.

**Table 4.** Performance comparison of our DeepBindGCN_RG_x with other methods in predicting experimental affinity on the PDBbind v.2016 core set (CASF-2016 core set) and v.2013 core set.

| Test set | Methods | Rmse | Pearson R | Spearman R |
| --- | --- | --- | --- | --- |
| PDBbind v.2016 core set | DeepBindGCN_RG_x | 1.3843 | 0.7719 | 0.7672 |
|  | KDEEP | 1.27 | 0.82 |  |
|  | Pafnucy | 1.42 | 0.78 |  |
|  | midlevel fusion | 1.308 | 0.810 | 0.807 |
|  | GraphBAR(dataset 4, Adj-2) | 1.413 | 0.778 |  |
|  | AK-score-ensemble | 1.293 |  |  |
|  | DeepAtom | 1.23 | 0.831 |  |
|  | PointNet(B) | 1.26 | 0.831 | 0.827 |
|  | PointTransform(B) | 1.19 | 0.859 | 0.853 |
|  | AEScore | 1.22 | 0.83 |  |
|  | ResAtom-Score |  | 0.833 |  |
|  | DEELIG |  | 0.889 |  |
|  | PIGNet (ensemble) |  | 0.761 |  |
|  | BAPA | 1.308 |  |  |
| PDBbind v.2013 core set | DeepBindGCN_RG_x | 1.4864 | 0.7503 | 0.7358 |
|  | SE-OnionNet | 1.692 | 0.812 |  |
|  | DeepBindRG | 1.817 | 0.6394 |  |
|  | DEELIG |  | 0.894 |  |
|  | GraphBAR(dataset4, best) | 1.636 | 0.704 |  |
|  | BAPA | 1.457 |  |  |

To explore whether the vector representation of the amino acid has a better performance than the onehot representation, we have trained a model with onehot representation with the same model architecture and training and validation set. The performance over validation with different epochs is shown in Figure S10 and Table S6 and S7. We can observe that its performance is not good as DeepBindGCN.

Like the DFCNN(Zhang, Lin, *et al*., 2022; H. Zhang, Liao, Cai, *et al*., 2019), the DeepBindGCN can be applied to quickly and accurately identify the potential protein target. The DeepBindGCN has inherited the efficiency of the DFCNN model, which is also not dependent on protein-ligand docking structure. In the meantime, the DeepBindGCN is much more efficient in keeping the spatial information within ligands and pockets through graphic representation. Since spatial information is critical in many protein-ligand interactions, the DeepBindGCN should be more useful in target identification for given compounds through inverse target searching.

Also, similar to DFCNN or autodock vina(Trott and Olson, 2010) being applied in our previous work for specificity estimation of a given compound(Zhang, Gong, *et al*., 2022), the DeepBindGCN can also be used to calculate the specificity similarly. Our proposed scoring is shown in Figure 6. To estimate the specificity for large amounts of compounds, we can first use the DeepBindGCN_BC to make the reverse prediction against 102 proteins from DUD.E. We have defined a function to estimate the DeepBindGCN_BC-based specificity. The formula is used as follows:

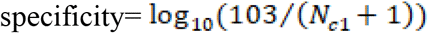

Where *N_c1_* is the counted number of compounds that have a DeepBindGCN_BC score larger than 0.9 during the reverse DFCNN prediction with 102 protein targets.

**Figure 6.**
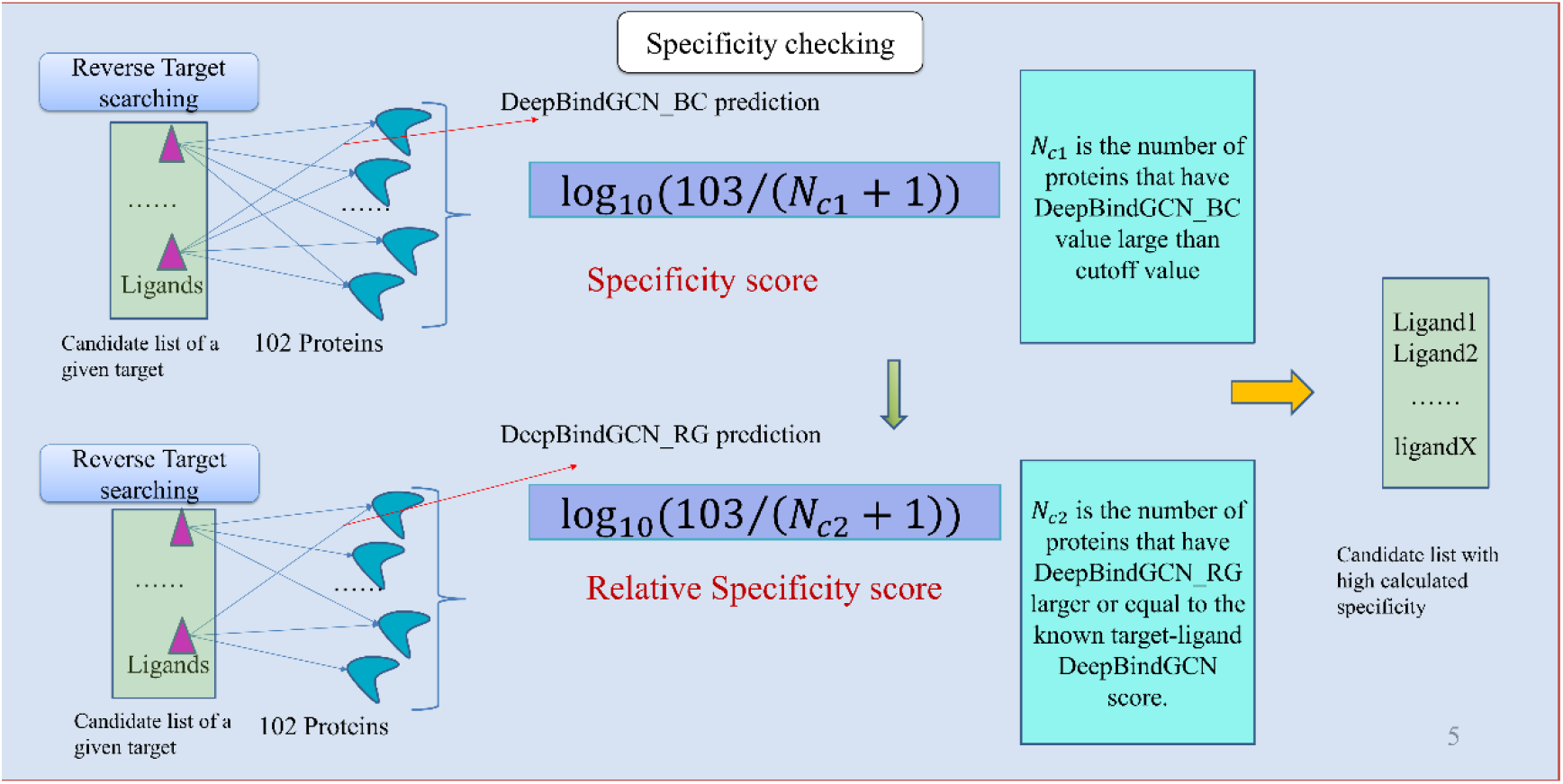
Our proposed specificity calculation strategy for virtual screening.

However, DeepBindGCN_BC doesn’t consider the binding affinity with these off-target, hence we can carry a DeepBindGCN_RG for further relative specificity. The relative specificity is calculated by following the formulas.

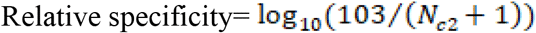

Where the *N_c2_* is the counted number of proteins that have a DeepBindGCN_RG score smaller than the known target-ligand DeepBindGCN_RG score. For instance, if we estimate the specificity of candidate compound G858-0261 of TIPE3, the *N_c2_* is the counted number of proteins (belonging to 102 targets from DUD.E) that have DeepBindGCN_RG score smaller than TIPE3-Y020-0019’s score 9.0349.

There is still space for improvement of the model in the future. We can test other model architectures, such as ATN, instead of GCN. We can apply molecular vectors in compounds as well. For instance, each chemical group was represented as a node with its molecular vector, and the edge was defined as chemical group neighbors. Also, we can add compounds molecular vectors as independent input. Furthermore, we can also integrate protein-ligand interaction pair information as graphic input, just as Moesser *et al.* has done(Moesser *et al*., 2022). Moreover, a similar strategy can be applied to protein-protein or protein-peptide interaction prediction. The protein interaction interface can be represented by graphic representation in a very similar way. Hence our work can provide helpful insight into protein-protein interaction or protein-peptide interaction prediction.

## Conclusion

We have developed DeepBindGCN_BC to identify accurate protein-ligand binding, and DeepBindGCN_RG to further estimate the protein-ligand binding affinity. Our GCN-based model not only help to identify binding ligands but also help to identify strong binding ligands, which are often more likely to be developed into drugs. The models have taken advantage of the graphic convolution network to represent spatial information efficiently. Also, we have added the molecular vector representation to enhance the pocket physical-chemical feature. Furthermore, we have tested the model in a much diversified DUD.E dataset and achieved good performance, indicating the reliability and practicality of our method. Also, to demonstrate its application in virtual screening, we have developed a pipeline and screened it over three cancer-related therapeutic targets, TIPE3 and PD-L1 dimer, as proof-of-concept applications. We also highlight its potential in other tasks, such as inverse target screening, specificity calculation, and iteratively screening *de novo* compounds by integrating with molecule generative models. We have deposited the source codes of our model on GitHub for user’s convenience. The models and the screening pipeline presented here would greatly help to facilitate computer-aided drug development.

## Key Points

- The present work demonstrates that the resulting model is accurate and extremely fast by using GCN and molecular vectors to represent the protein pocket effectively and compounds spatial information and physico-chemical.
- We have developed a binary classifier model that includes negative data during training to identify whether compounds will bind to a given target. Also, we have developed an affinity prediction model, which can further identify high-affinity binding compounds from the candidate list predicted by the binary classifier model.
- The developed DeepBindGCN model is a generalized protein-ligand prediction model, which is suitable for application to a wide range of therapeutic targets. In this work, we have applied DeepBindGCN on virtual screening against TIPE3 and PD-L1 dimer as proof-of-concept examples. The obtained candidate lists would help drug development against this target.

## Availability of data and materials

The proposed models and the scripts are available in GitHub public repositories (https://github.com/haiping1010/DeepBindGCN).

## Author contributions

HZ and JZ designed the study. HZ, KMS performed computations and data analyses. All authors contributed to writing the manuscript. HZ, and JZ supervised the study. All authors read and approved the final manuscript.

## Competing Interests

No authors have a conflict of interest in publishing this paper.

## Acknowledgments

This study was supported in part by the National Science Foundation of China (Grant No. 62106253, 21933010, 22250710136).

## Notes

### Competing Interest Statement

The authors have declared no competing interest.

## Reference

Ahmed, A. et al. (2021) DEELIG: A Deep Learning Approach to Predict Protein-igand Binding Affinity. Bioinform. Biol. Insights, 15, 11779322211030364.

Chen, J. et al. (2021) Chemical toxicity prediction based on semi-supervised learning and graph convolutional neural network. J. Cheminform.

Chen, Z. et al. (2021) ILearnPlus: A comprehensive and automated machine-learning platform for nucleic acid and protein sequence analysis, prediction and visualization. Nucleic Acids Res.

Fayngerts, S.A. et al. (2014) TIPE3 is the transfer protein of lipid second messengers that promote cancer. Cancer Cell.

Guzik, K. et al. (2017) Small-Molecule Inhibitors of the Programmed Cell Death-1/Programmed Death-Ligand 1 (PD-1/PD-L1) Interaction via Transiently Induced Protein States and Dimerization of PD-L1. J. Med. Chem.

Humphrey, W. et al. (1996) VMD: visual molecular dynamics. J. Mol. Graph., 14, 33-8, 27–8.

Jiménez, J. et al. (2018) KDEEP: Protein-Ligand Absolute Binding Affinity Prediction via 3D-Convolutional Neural Networks. J. Chem. Inf. Model.

Jones, D. et al. (2021) Improved Protein–Ligand Binding Affinity Prediction with Structure-Based Deep Fusion Inference. J. Chem. Inf. Model., 61, 1583–1592.

Klebe, G. (2013) Protein-Ligand Interactions as the Basis for Drug Action. In, Drug Design.

Kojima, R. et al. (2020) KGCN: A graph-based deep learning framework for chemical structures. J. Cheminform.

Kwon, Y. et al. (2020) AK-Score: Accurate Protein-Ligand Binding Affinity Prediction Using an Ensemble of 3D-Convolutional Neural Networks. Int. J. Mol. Sci., 21.

Landrum, G. (2006) RDKit: Open-source Cheminformatics. Http://Www.Rdkit.Org/.

Li, Q. et al. (2021) TIPE3 promotes non-small cell lung cancer progression via the protein kinase B/extracellular signal-regulated kinase 1/2-glycogen synthase kinase 3β-β-catenin/Snail axis. Transl. Lung Cancer Res.

Li, Y. et al. (2019) DeepAtom: A Framework for Protein-Ligand Binding Affinity Prediction. In, 2019 IEEE International Conference on Bioinformatics and Biomedicine (BIBM)., pp. 303–310.

Meli, R. et al. (2021) Learning protein-ligand binding affinity with atomic environment vectors. J. Cheminform., 13, 59.

Moesser, M.A. et al. (2022) Protein-Ligand Interaction Graphs: Learning from Ligand-Shaped 3D Interaction Graphs to Improve Binding Affinity Prediction. bioRxiv.

Moon, S. et al. (2022) PIGNet: a physics-informed deep learning model toward generalized drug–target interaction predictions. Chem. Sci., 13, 3661–3673.

Murtagh, F. and Contreras, P. (2012) Algorithms for hierarchical clustering: An overview. Wiley Interdiscip. Rev. Data Min. Knowl. Discov.

Mysinger, M.M. et al. (2012) Directory of Useful Decoys, Enhanced (DUD-E): Better Ligands and Decoys for Better Benchmarking. J. Med. Chem., 55, 6582–6594.

Nguyen, Thin et al. (2021) GraphDTA: Predicting drug target binding affinity with graph neural networks. Bioinformatics.

Pettersen, E.F. et al. (2004) UCSF Chimera - A visualization system for exploratory research and analysis. J. Comput. Chem.

Roy, A. et al. (2012) COFACTOR: an accurate comparative algorithm for structurebased protein function annotation. Nucleic Acids Res., 40, W471–7.

Savojardo, C. et al. (2018) DeepSig: Deep learning improves signal peptide detection in proteins. Bioinformatics.

Seo, S. et al. (2021) Binding affinity prediction for protein–ligand complex using deep attention mechanism based on intermolecular interactions. BMC Bioinformatics.

Son, J. and Kim, D. (2021) Development of a graph convolutional neural network model for efficient prediction of protein-ligand binding affinities. PLoS One, 16, e0249404.

Stepniewska-Dziubinska, M.M. et al. (2018) Development and evaluation of a deep learning model for protein-ligand binding affinity prediction. Bioinformatics.

Torng, W. and Altman, R.B. (2019) Graph Convolutional Neural Networks for Predicting Drug-Target Interactions. J. Chem. Inf. Model.

Trott, O. and Olson, A.J. (2010) AutoDock Vina: improving the speed and accuracy of docking with a new scoring function, efficient optimization, and multithreading. J. Comput. Chem., 31, 455–61.

Visualizer, D.S. (2005) v4. 0.100. 13345. Accelrys Softw. Inc.

Wang, S. et al. (2021) SE-OnionNet: A Convolution Neural Network for Protein-Ligand Binding Affinity Prediction. Front. Genet., 11.

Wang, Y. et al. (2022) A point cloud-based deep learning strategy for protein-ligand binding affinity prediction. Brief. Bioinform., 23, bbab474.

Wang, Y. et al. (2021) ResAtom System: Protein and Ligand Affinity Prediction Model Based on Deep Learning.

Yuan, H. et al. (2021) Protein-ligand binding affinity prediction model based on graph attention network. Math. Biosci. Eng.

Zhang, H., Gong, X., et al. (2022) An Efficient Modern Strategy to Screen Drug Candidates Targeting RdRp of SARS-CoV-2 With Potentially High Selectivity and Specificity. Front. Chem., 10.

Zhang, H., Li, J., et al. (2021) An Integrated Deep Learning and Molecular Dynamics Simulation-Based Screening Pipeline Identifies Inhibitors of a New Cancer Drug Target TIPE2. Front. Pharmacol., 12, 3297.

Zhang, H., Liao, L., Saravanan, K.M., et al. (2019) DeepBindRG: a deep learning based method for estimating effective protein-ligand affinity. PeerJ, 7, e7362.

Zhang, H., Saravanan, K.M., et al. (2022) Generating and screening de novo compounds against given targets using ultrafast deep learning models as core components. Brief. Bioinform., bbac226.

Zhang, H., Liao, L., Cai, Y., et al. (2019) IVS2vec: A tool of Inverse Virtual Screening based on word2vec and deep learning techniques. Methods, 166, 57–65.

Zhang, Haiping, Zhang, T., et al. (2021) A novel virtual drug screening pipeline with deep-leaning as core component identifies inhibitor of pancreatic alpha-amylase. In, Proceedings - 2021 IEEE International Conference on Bioinformatics and Biomedicine, BIBM 2021.

Zhang, Haiping et al. (2020) A novel virtual screening procedure identifies Pralatrexate as inhibitor of SARS-CoV-2 RdRp and it reduces viral replication in vitro. PLoS Comput. Biol., 16, e1008489.

Zhang, Haiping, Zhang, T., et al. (2022) DeepBindBC: A practical deep learning method for identifying native-like protein-ligand complexes in virtual screening. Methods, 205, 247–262.

Zhang, Haiping, Lin, X., et al. (2022) Validation of Deep Learning-Based DFCNN in Extremely Large-Scale Virtual Screening and Application in Trypsin I Protease Inhibitor Discovery. Front. Mol. Biosci., 9.

Zhang, S. et al. (2019) Graph convolutional networks: a comprehensive review. Comput. Soc. Networks.

Zhao, Q. et al. (2019) AttentionDTA: Prediction of drug-target binding affinity using attention model. In, Proceedings - 2019 IEEE International Conference on Bioinformatics and Biomedicine, BIBM 2019.

