## Supplementary materials for "DeepBindGCN: Integrating Molecular Vector Representation with Graph Convolutional Neural Networks for Accurate Protein-Ligand Interaction Prediction"

**Supplementary material section 1:**

To obtain the dataset for DeepBindGCN_RG_x, we excluded PDBbind v.2016 core set (CASF-2016 core set) and v.2013 core set from PDBbind database, and finally obtained 16,584 protein-ligand pairs. The dataset was divided into training data set and validation set, which contains 16,000 and 584 samples, respectively. The v.2013 core set (195 protein-ligand pairs) and PDBbind v.2016 core set (285 protein-ligand pairs) were kept as testing set. The architecture is same as the DeepBindGCN_RG and the total training epoch is 2000, the models were saved every 100 epochs.

**Supplementary material section 2:**

**Detailed procedure of MD, pocket MD and metadynamics simulation**

The initial protein-compound complexes were from the top score conformation Schrödinger docking, the ligand was edited by pymol software [1] to make it in correct protonation state at pH 7.

We performed MD simulation for the TIPE3-compound complexes. Notably, we have cutted “FSSKSLALQAQKKILSKIAS” (amino acid 110 to 129) part of the TIPE3 to save the simulation resource. To save the computational resources, we have carried pocket MD for DNA-PKcs-compound and PD-L1-compound complexes by only keeping the binding pocket region for simulation. Binding free energy calculation can be estimated by metadynamics simulations to explore whether protein-ligand will bind in solution. Metadynamics relies on addition of a bias potential to sample the free energy landscape along a specific collective variable of interest [2,3]. Note that the binding free energy calculations from Metadynamics may only be suitable for detect the general trend of binding in virtual screening.

The pocket MD is same as the classical MD simulation, except that we only using the pocket region to reduce system size for simulation [4], which is inspired by a previous dynamic undocking (DUck) method [5]. An in-house script was used to extract the pocket region of the protein (here, we used 1.2nm within the binding ligand), the N terminal and C terminal ends were capped with the ACE and NHE terminals, respectively. We applied position restrains to the ACE and NHE terminals to maintain the relative conformation of the pocket. MD simulation or pocket MD simulation was carried out by Gromacs with AMBER-99SB force field [6,7]. The topology of ligand and the partial charges of ligand was generated by ACPYPE [8], which relies on Antechamber [9]. Firstly, we created a dodecahedron box and put the target-ligand complex at the center. A minimum distance from the protein to box edge was set to 1 nm. We filled the dodecahedron box with TIP3P water molecules [10], the counter ions were added to neutralize the total charge using the Gromacs program tool [11]. The long-range electrostatic interactions under the periodic boundary conditions was calculated with Particle Mesh Ewald approach [12]. A cutoff of 14 Å was used for van der Waals non-bonded interactions. Covalent bonds involving hydrogen atoms were constrained by applying the LINCS algorithm [13].

We performed the energy minimization steps with a step-size of 0.001ns, 100 ps simulation with isothermal-isovolumetric ensemble (NVT), and 10ns simulation with isothermal-isobaric ensemble (NPT) for water equilibrium. After that, a 40ns NPT production run (step size 2 fs) was carried out. The Parrinello-Rahman barostat and the modified Berendsen thermostat were used for simulation with a fixed temperature of 308 K and a pressure of 1 atm. RMSD and hydrogen bond number of the trajectory were calculated using Gromacs tools.

The simulation was continued using the metadynamics approach for exploring the free energy landscape. We carried 40ns metadynamics simulation with Plumed[14] patched Gromacs. The interface coordination number of atoms of protein ligand complex was used as collective variable (CV). The protein-ligand interface coordination numbers correlate with the numbers of atom contact, and larger coordination number usually indicates that protein-ligand is in binding state.

The coordination number $C$ is defined as follows by Plumed:

$$C = \sum_{i \in A} \sum_{j \in B} S_{ij} \quad (1) \quad \text{and}$$

$$S_{ij} = \frac{1 - \left( \frac{r_{ij} - d_0}{r_0} \right)^n}{1 - \left( \frac{r_{ij} - d_0}{r_0} \right)^m} \quad (2)$$

In the simulation, $n$ was 6, $m$ was 12, $d_0$ was 0 nm and $r_0$ was 0.5 nm. $d_0$ is a parameter of the switching function. $r_{ij}$ is the distance between atom $i$ and atom $j$. The degrees of contacts between two groups of atoms can be estimated by above function(1) [14]. Metadynamics simulation for each protein-ligand system was performed for 40 ns. During the metadynamics simulation, Gaussian values were deposited every 1 ps with a height of 0.3 kJ/mol. The widths of the Gaussians were 5 for the coordination number. The free energy landscapes of the metadynamics simulations along the CV were generated by the Plumed program and plotted using Gnuplot [15].

### Supplementary Figures:

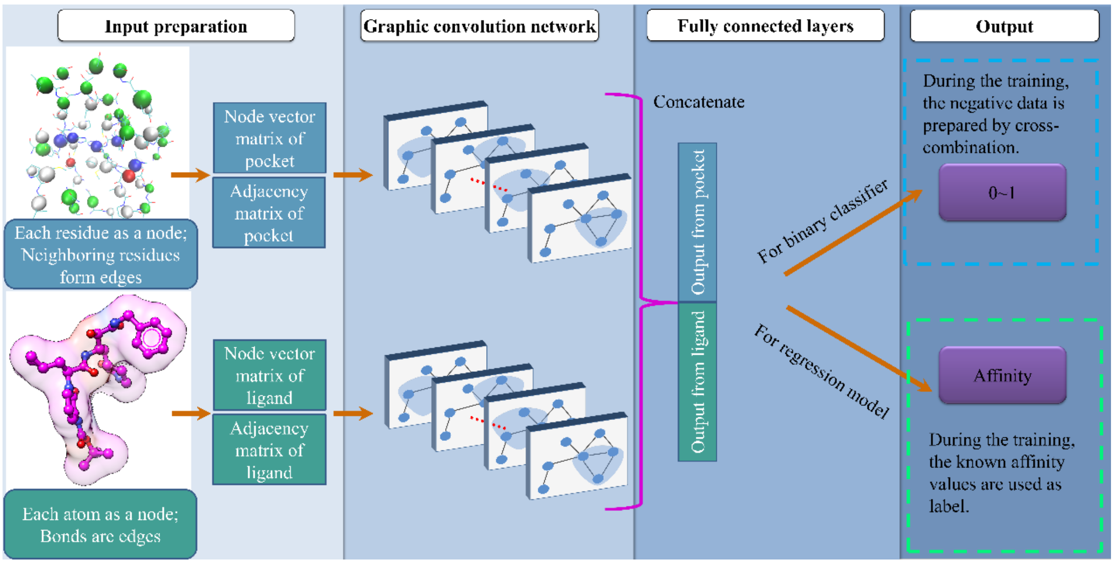


**Figure S1. Illustration of the model constructing flow.**

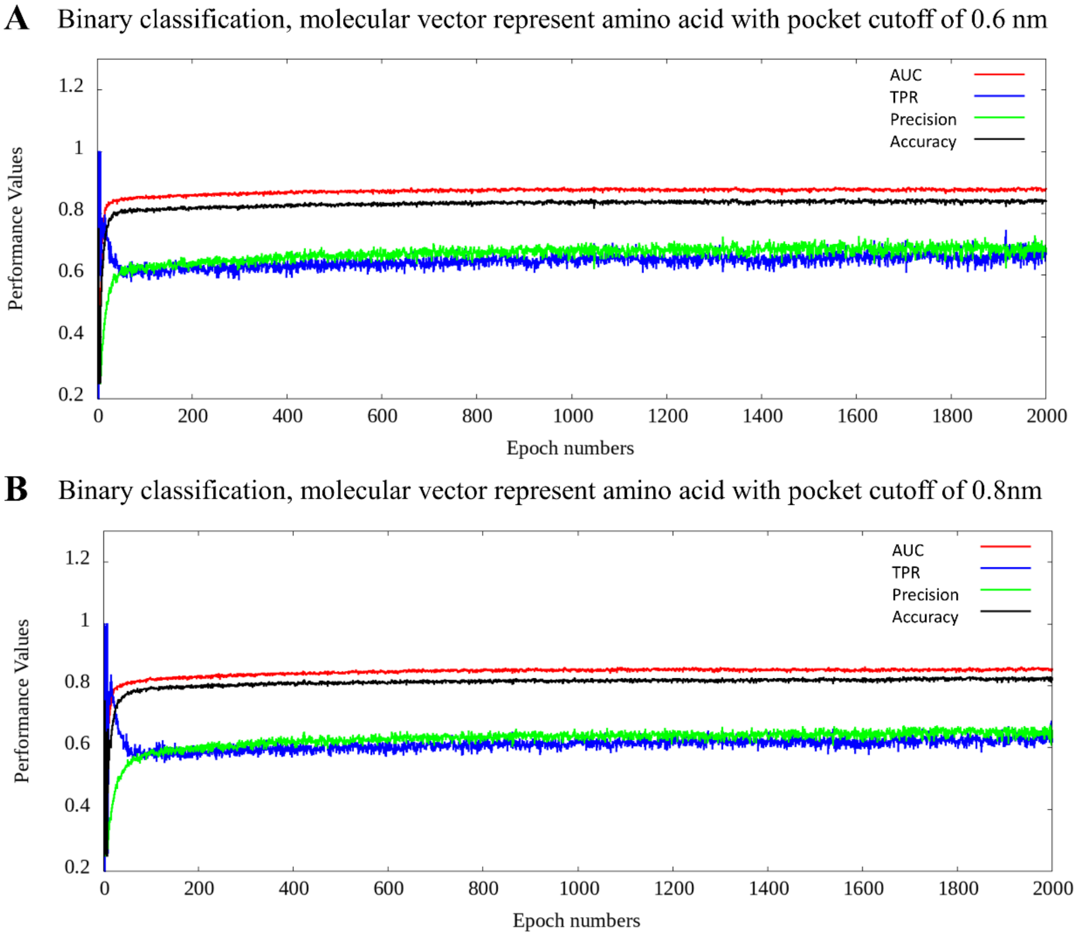


**Figure S2.** The DeepBindGCN_BC performance over the 2000 epoch training. A.

pocket cutoff using 0.6nm, B, pocket cutoff using 0.8nm. The amino acid in the pocket is represented as molecular vector.

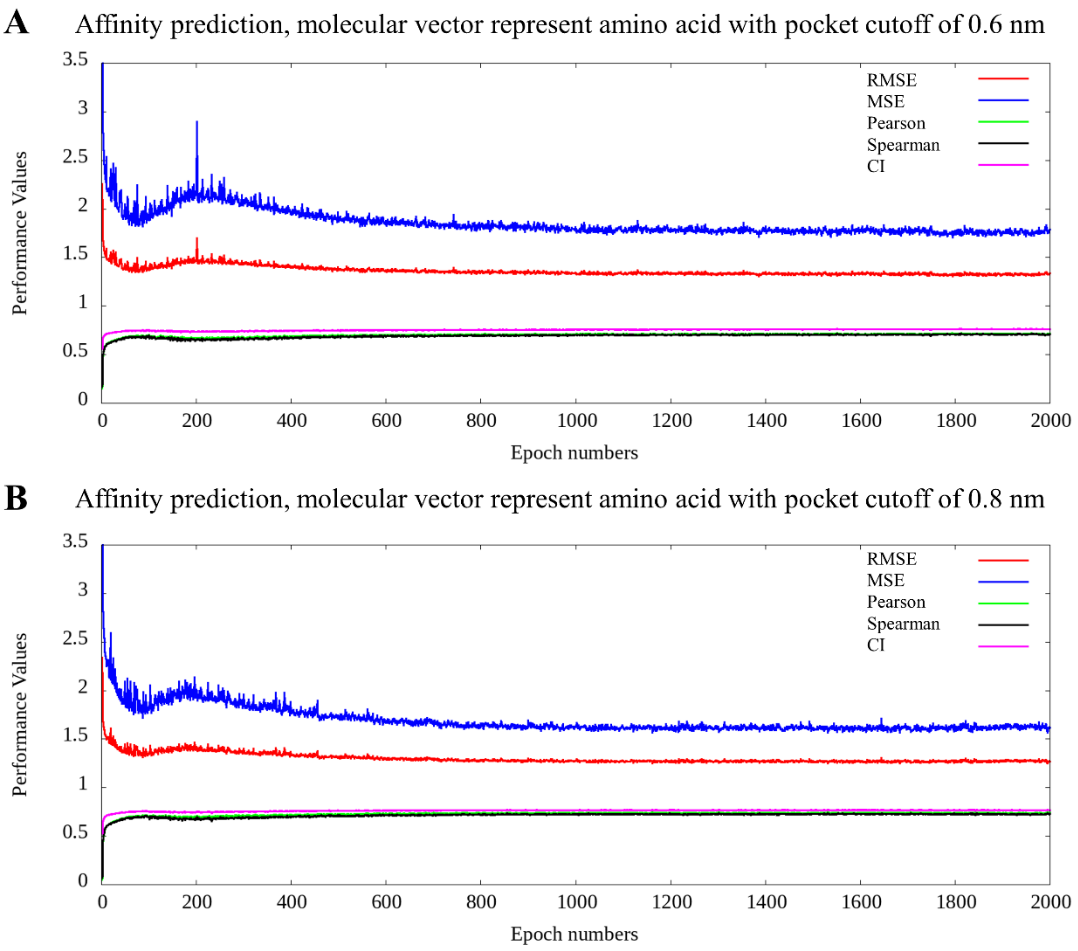


**Figure S3. The DeepBindGCN_RG performance over the 2000 epoch training.** A. pocket cutoff using 0.6nm, B, pocket cutoff using 0.8nm. The amino acids in the pocket are represented as molecular vectors.

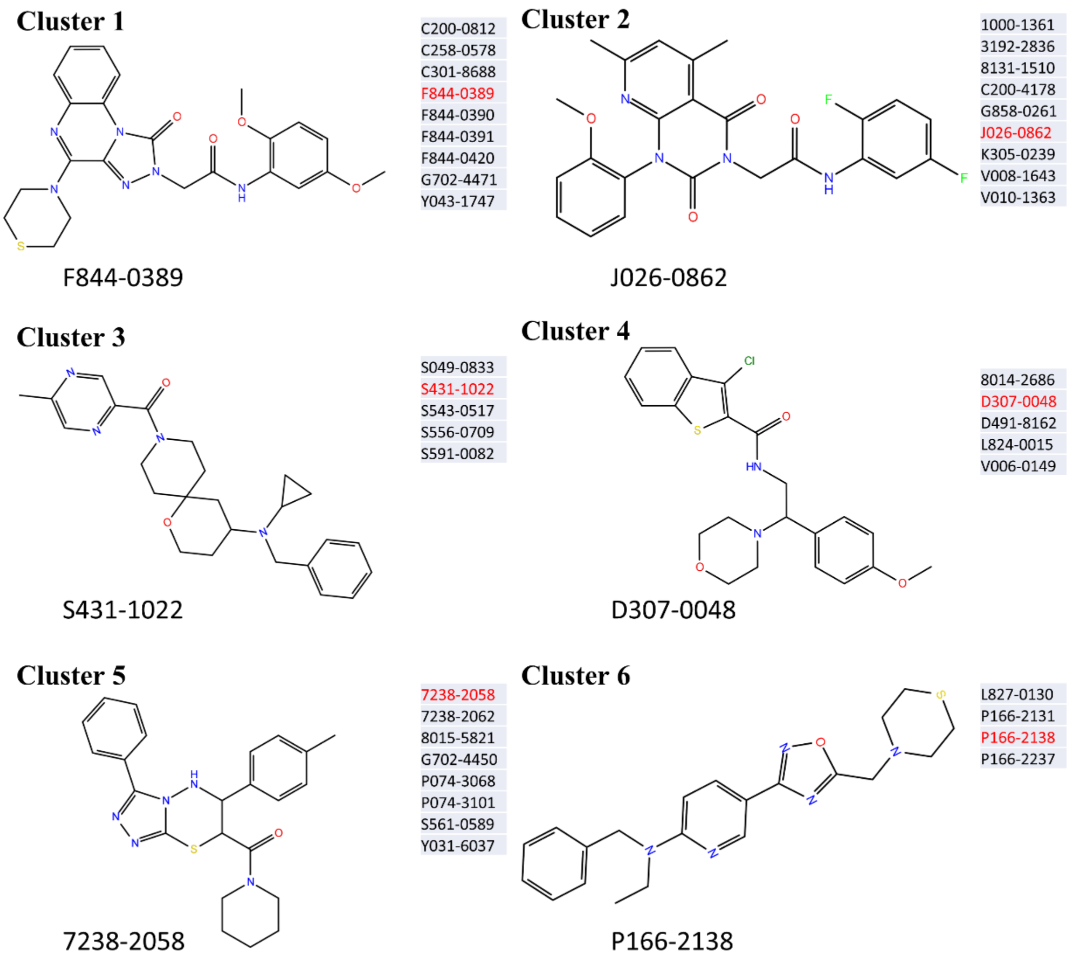


**Figure S4. The six clusters and corresponding representative cluster center structure of top predicted candidates from DeepBindGCN_BC and DeepBindGCN_RG for the TIPE3.**

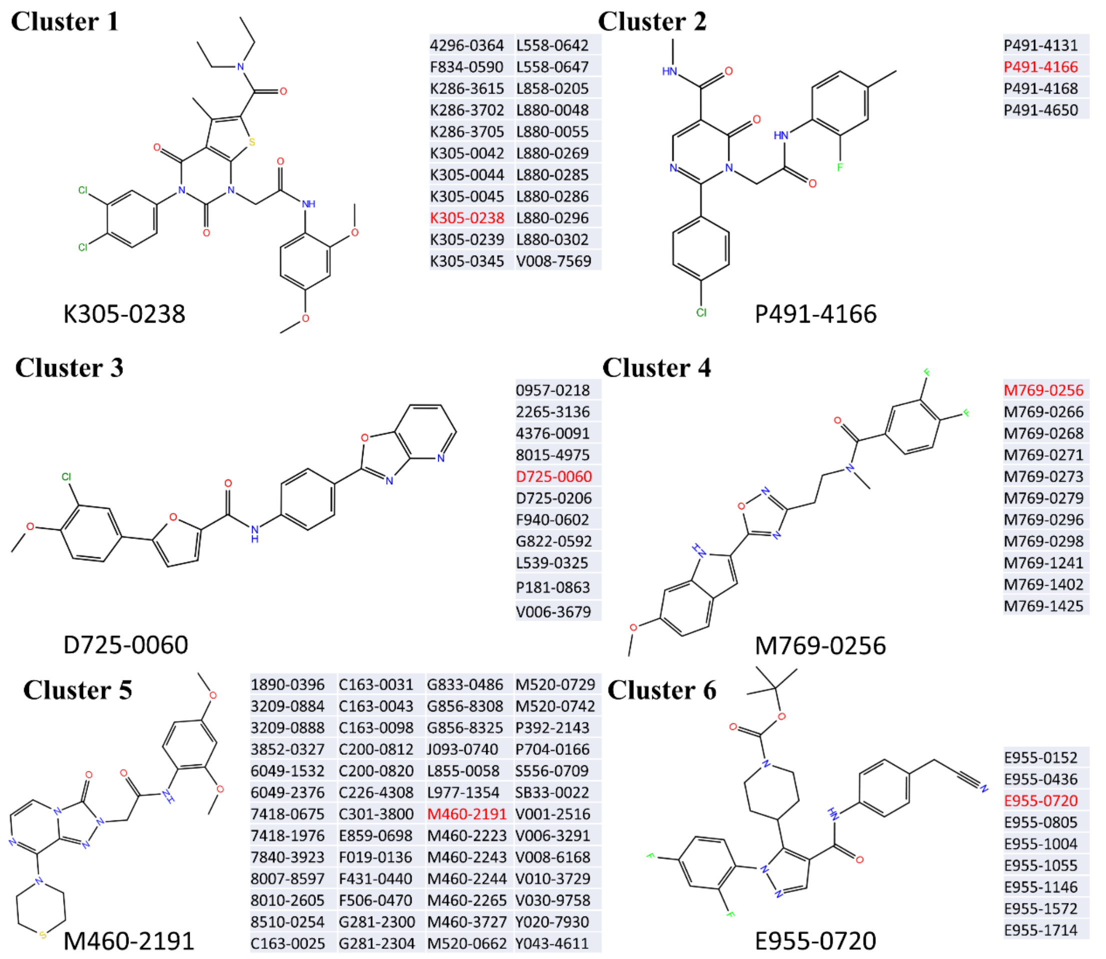


**Figure S5. The six clusters and corresponding representative cluster center structure of top predicted candidates from DeepBindGCN_BC and DeepBindGCN_RG for the PD-L1.**

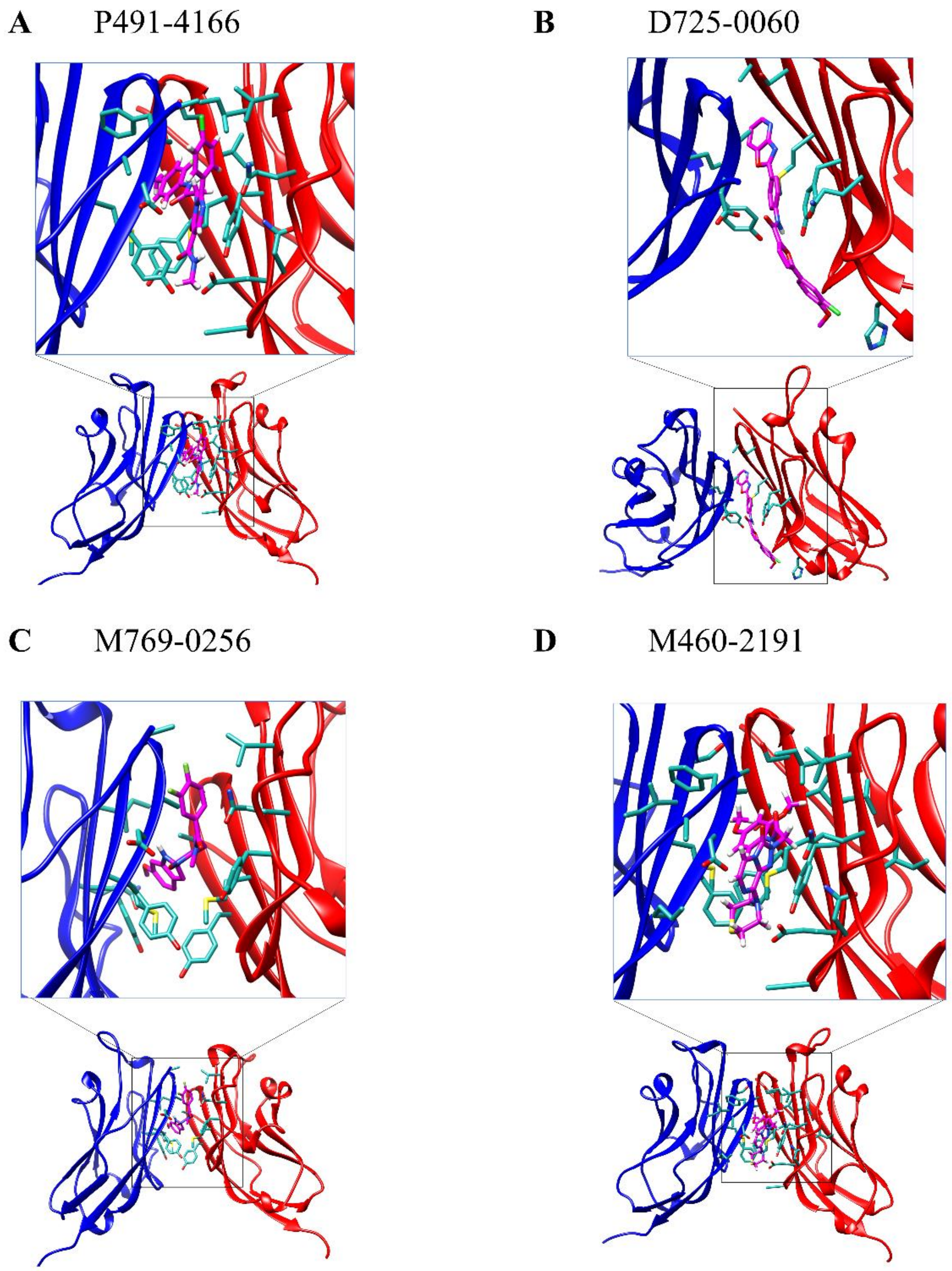


**Figure S6. The snapshot and 2D plot of PD-L1 with representative cluster center compounds from docking.**

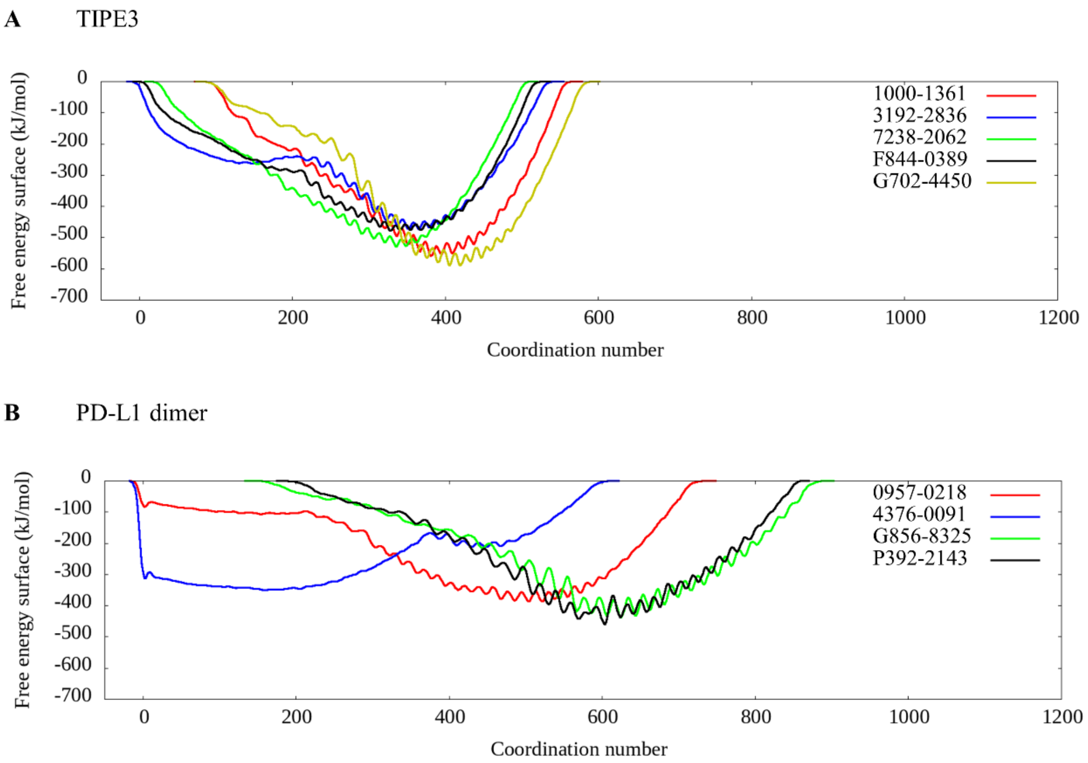


**Figure S7. The calculated free energy landscape from metadynamics simulation for those candidates that have favorable binding with the given target.** A, The calculated free energy landscape for TIPE3 with selected candidates; B, The calculated free energy landscape for PD-L1 with selected candidates.

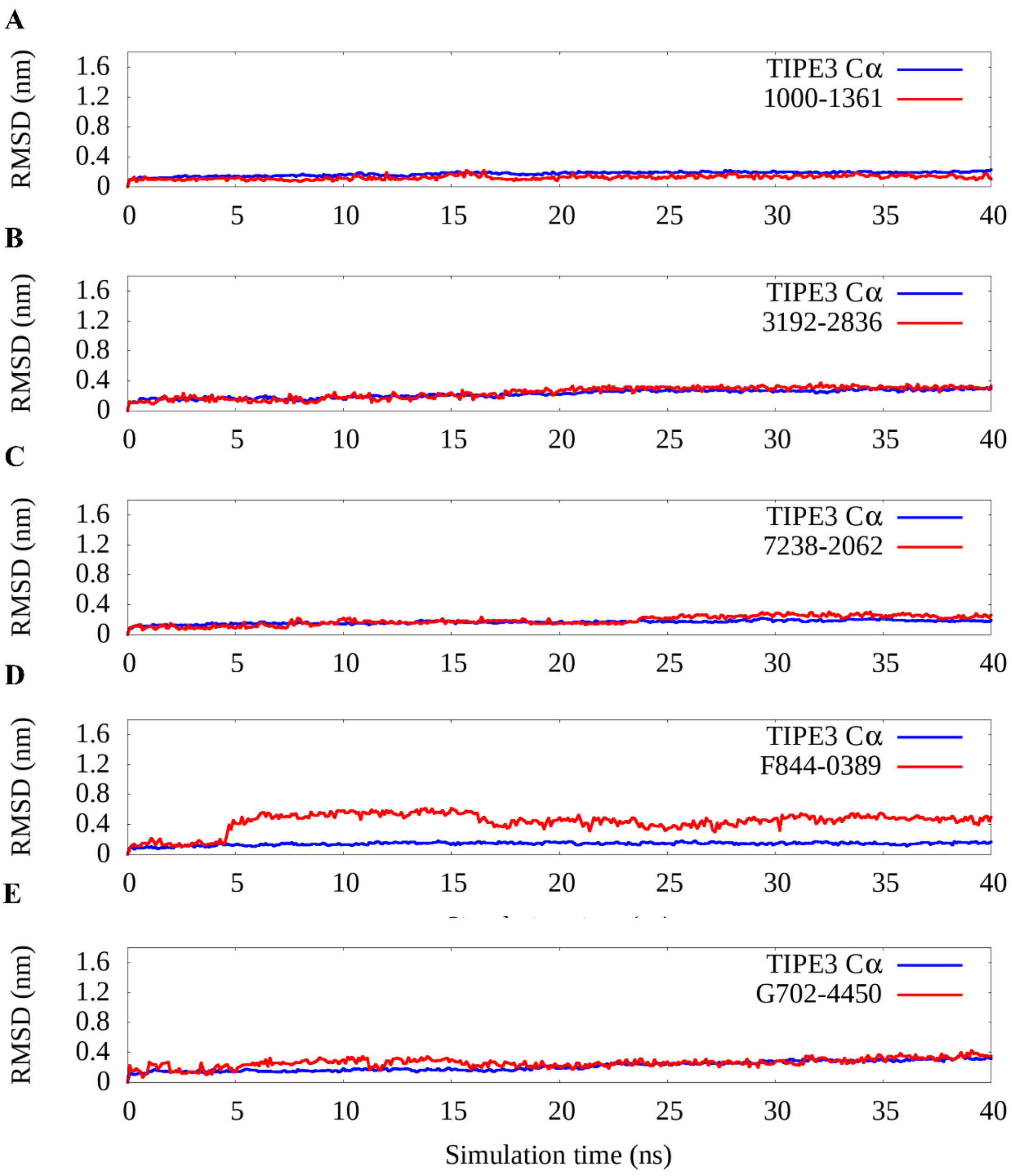


**Figure S8. The rmsd value of selected compounds with the TIPE3 during the MD simulation.**

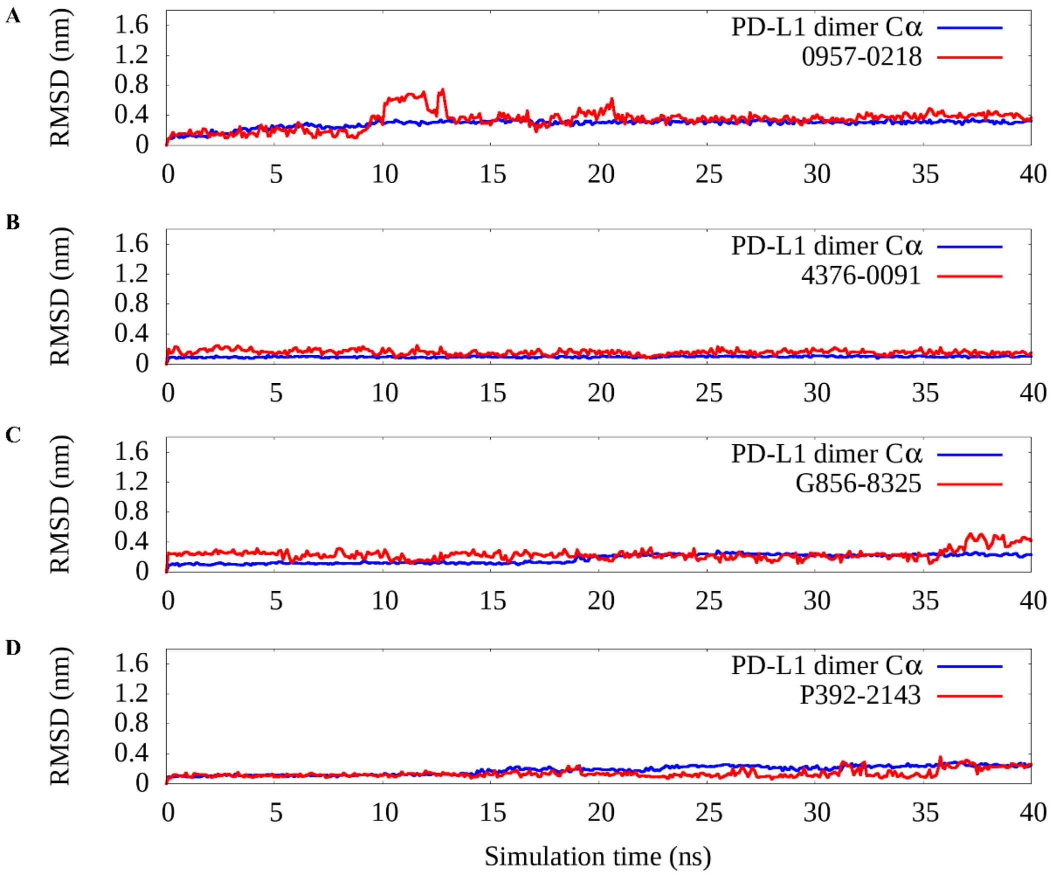


**Figure S9. The rmsd value of selected compounds with the PD-L1 during the MD simulation.**

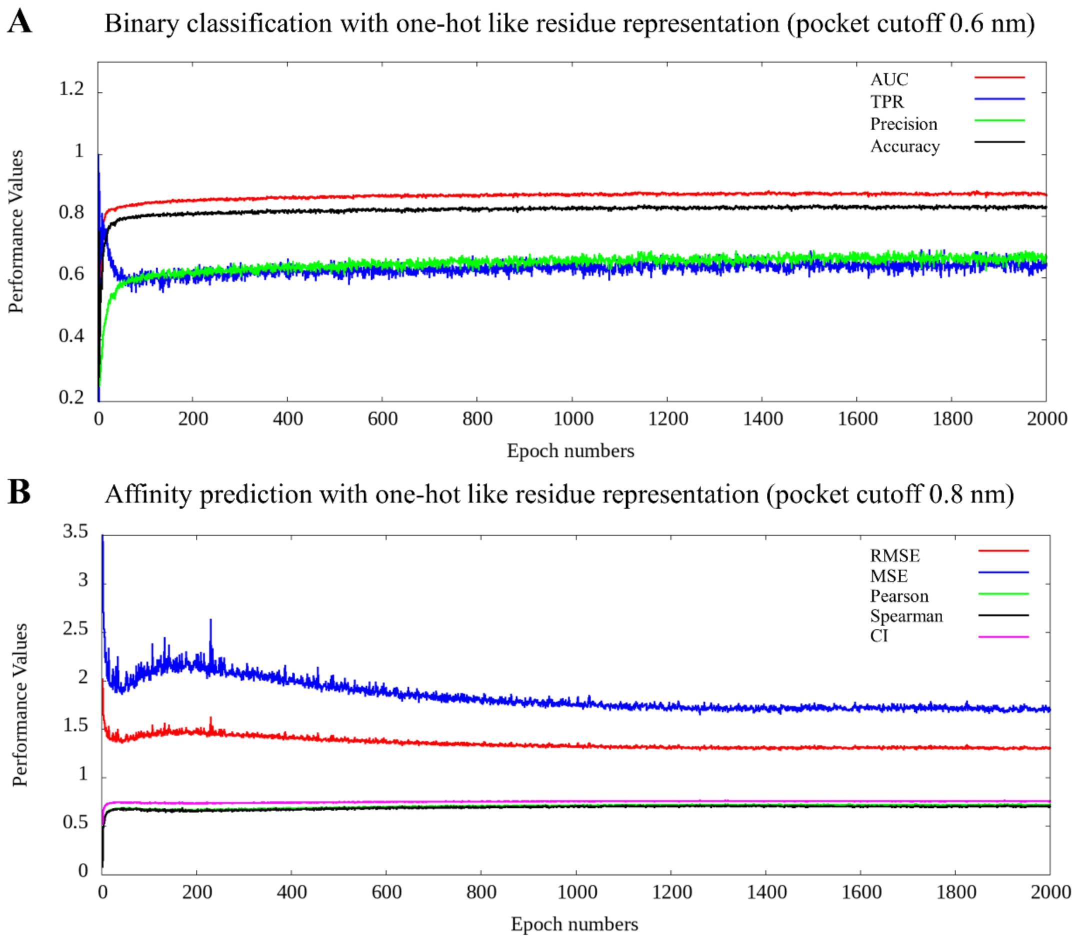


**Figure S10. The performance of binary classification and affinity prediction models with one-hot like residue representation over the 2000 epoch training.** A. pocket cutoff using 0.6nm, B, pocket cutoff using 0.8nm. The amino acid in the pocket is represented as one-hot representation.

### Supplementary Tables:

**Table S1. We listed performance of DeepBindGCN_BC on test set during the training with epoch interval 100.**

| Pocket cutoff | Epoch | AUC | TPR | Precision | Accuracy |
|---|---|---|---|---|---|
| 0.6nm | 100 | 0.8493 | 0.6408 | 0.6096 | 0.8076 |
| | 200 | 0.8589 | 0.6138 | 0.6418 | 0.8178 |
| | 300 | 0.8626 | 0.6508 | 0.6246 | 0.8149 |
| | 400 | 0.8705 | 0.6254 | 0.6737 | 0.8306 |
| | 500 | 0.8676 | 0.6367 | 0.6524 | 0.8244 |
| | 600 | 0.8682 | 0.6350 | 0.6626 | 0.8279 |
| | 700 | 0.8750 | 0.6233 | 0.6831 | 0.8335 |
| | 800 | 0.8745 | 0.6508 | 0.6588 | 0.8284 |

| | | | | | |
|---|---|---|---|---|---|
| | 900 | 0.8789 | 0.6296 | 0.6837 | 0.8346 |
| | 1000 | 0.8767 | 0.6400 | 0.6802 | 0.8348 |
| | 1100 | 0.8703 | 0.6438 | 0.6747 | 0.8333 |
| | 1200 | 0.8744 | 0.6483 | 0.6801 | 0.8358 |
| | 1300 | 0.8719 | 0.6704 | 0.6632 | 0.8325 |
| | 1400 | 0.8755 | 0.6579 | 0.6765 | 0.8358 |
| | 1500 | 0.8790 | 0.6338 | 0.7071 | 0.8428 |
| | 1600 | 0.8752 | 0.6488 | 0.6929 | 0.8403 |
| | 1700 | 0.8775 | 0.6625 | 0.6780 | 0.8370 |
| | 1800 | 0.8738 | 0.6567 | 0.6915 | 0.8409 |
| | 1900 | 0.8778 | 0.6667 | 0.6917 | 0.8424 |
| | 2000 | 0.8788 | 0.6863 | 0.6767 | 0.8396 |
| | 100 | 0.8222 | 0.5933 | 0.5901 | 0.7953 |
| | 200 | 0.8275 | 0.6021 | 0.5879 | 0.7950 |
| | 300 | 0.8367 | 0.5988 | 0.6186 | 0.8074 |
| | 400 | 0.8401 | 0.5875 | 0.6242 | 0.8084 |
| | 500 | 0.8403 | 0.6013 | 0.6304 | 0.8122 |
| | 600 | 0.8454 | 0.5904 | 0.6326 | 0.8119 |
| | 700 | 0.8443 | 0.6138 | 0.6236 | 0.8108 |
| | 800 | 0.8493 | 0.6154 | 0.6280 | 0.8127 |
| | 900 | 0.8513 | 0.6142 | 0.6420 | 0.8179 |
| 0.8nm | 1000 | 0.8493 | 0.6154 | 0.6422 | 0.8181 |
| | 1100 | 0.8502 | 0.6171 | 0.6329 | 0.8148 |
| | 1200 | 0.8559 | 0.6088 | 0.6543 | 0.8218 |
| | 1300 | 0.8560 | 0.6250 | 0.6394 | 0.8181 |
| | 1400 | 0.8492 | 0.6300 | 0.6340 | 0.8166 |
| | 1500 | 0.8539 | 0.6079 | 0.6479 | 0.8194 |
| | 1600 | 0.8550 | 0.6238 | 0.6531 | 0.8231 |
| | 1700 | 0.8517 | 0.6017 | 0.6600 | 0.8229 |
| | 1800 | 0.8489 | 0.6154 | 0.6527 | 0.8220 |
| | 1900 | 0.8546 | 0.6042 | 0.6648 | 0.8249 |
| | 2000 | 0.8537 | 0.6175 | 0.6552 | 0.8231 |

**Table S2. We listed performance of DeepBindGCN_RG on test set during the training with epoch interval 100.**

| | Epoch | Rmse: | Mse | Pearson | Spearman | CI |
|---|---|---|---|---|---|---|
| | 100 | 1.3717 | 1.8816 | 0.7040 | 0.6968 | 0.7550 |
| | 200 | 1.4645 | 2.1449 | 0.6675 | 0.6524 | 0.7373 |
| | 300 | 1.4335 | 2.0550 | 0.6813 | 0.6684 | 0.7437 |
| | 400 | 1.4051 | 1.9744 | 0.6875 | 0.6725 | 0.7456 |
| | 500 | 1.3681 | 1.8716 | 0.6961 | 0.6860 | 0.7509 |
| | 600 | 1.3574 | 1.8426 | 0.7025 | 0.6915 | 0.7536 |

| | | | | | |
|---|---|---|---|---|---|
| 700 | 1.3526 | 1.8296 | 0.7043 | 0.6927 | 0.7540 |
| 800 | 1.3489 | 1.8195 | 0.7042 | 0.6949 | 0.7553 |
| 900 | 1.3681 | 1.8717 | 0.7095 | 0.7003 | 0.7572 |
| 1000 | 1.3308 | 1.7709 | 0.7108 | 0.7034 | 0.7591 |
| 1100 | 1.3293 | 1.7670 | 0.7114 | 0.7038 | 0.7592 |
| 1200 | 1.3417 | 1.8001 | 0.7084 | 0.7025 | 0.7585 |
| 1300 | 1.3277 | 1.7628 | 0.7103 | 0.7025 | 0.7585 |
| 1400 | 1.3483 | 1.8179 | 0.7067 | 0.6991 | 0.7573 |
| 1500 | 1.3310 | 1.7715 | 0.7096 | 0.7030 | 0.7595 |
| 1600 | 1.3291 | 1.7666 | 0.7118 | 0.7052 | 0.7599 |
| 1700 | 1.3318 | 1.7738 | 0.7111 | 0.7043 | 0.7603 |
| 1800 | 1.3327 | 1.7761 | 0.7106 | 0.7047 | 0.7602 |
| 1900 | 1.3277 | 1.7627 | 0.7138 | 0.7067 | 0.7609 |
| 2000 | 1.3361 | 1.7852 | 0.7141 | 0.7098 | 0.7628 |
| 100 | 1.3141 | 1.7270 | 0.7153 | 0.7137 | 0.7604 |
| 200 | 1.3331 | 1.7772 | 0.7192 | 0.7062 | 0.7572 |
| 300 | 1.2859 | 1.6534 | 0.7392 | 0.7256 | 0.7677 |
| 400 | 1.2966 | 1.6810 | 0.7276 | 0.7122 | 0.7620 |
| 500 | 1.2672 | 1.6059 | 0.7413 | 0.7311 | 0.7702 |
| 600 | 1.2465 | 1.5538 | 0.7497 | 0.7396 | 0.7752 |
| 700 | 1.2558 | 1.5770 | 0.7423 | 0.7309 | 0.7714 |
| 800 | 1.2201 | 1.4885 | 0.7580 | 0.7463 | 0.7779 |
| 900 | 1.2135 | 1.4725 | 0.7560 | 0.7498 | 0.7786 |
| 1000 | 1.2039 | 1.4494 | 0.7580 | 0.7500 | 0.7796 |
| 1100 | 1.2298 | 1.5124 | 0.7467 | 0.7393 | 0.7737 |
| 1200 | 1.1914 | 1.4195 | 0.7617 | 0.7475 | 0.7789 |
| 1300 | 1.1991 | 1.4378 | 0.7574 | 0.7448 | 0.7775 |
| 1400 | 1.2004 | 1.4410 | 0.7570 | 0.7446 | 0.7778 |
| 1500 | 1.2039 | 1.4494 | 0.7563 | 0.7435 | 0.7770 |
| 1600 | 1.2184 | 1.4844 | 0.7484 | 0.7370 | 0.7726 |
| 1700 | 1.2162 | 1.4791 | 0.7497 | 0.7400 | 0.7752 |
| 1800 | 1.2176 | 1.4825 | 0.7501 | 0.7372 | 0.7741 |
| 1900 | 1.2083 | 1.4599 | 0.7518 | 0.7398 | 0.7745 |
| 2000 | 1.2107 | 1.4657 | 0.7518 | 0.7410 | 0.7756 |

**Table S3. The performance of DeepBindGCN_BC (pocket cutoff 0.6nm) on the DUD.E dataset.**

| PDBID | AUC | TPR | Precision | Accuracy | MCC | data_size | pos_size | neg_size |
|---|---|---|---|---|---|---|---|---|
| 3BWM | 1.0000 | 0.8537 | 1.0000 | 0.8571 | 0.3492 | 42 | 41 | 1 |
| 1ZW5 | 0.5765 | 0.0118 | 1.0000 | 0.2500 | 0.0535 | 112 | 85 | 27 |
| 2AA2 | 0.7052 | 0.2217 | 1.0000 | 0.2290 | 0.0515 | 214 | 212 | 2 |
| 3KRJ | 0.9378 | 0.7558 | 1.0000 | 0.7589 | 0.1944 | 394 | 389 | 5 |

| | | | | | | | | |
|---|---|---|---|---|---|---|---|---|
| 3L3M | 0.8029 | 0.5301 | 1.0000 | 0.5355 | 0.1125 | 1057 | 1045 | 12 |
| 2OWB | 0.4205 | 0.0044 | 1.0000 | 0.1722 | 0.0273 | 273 | 227 | 46 |
| 3KBA | 0.1230 | 0.3615 | 0.9975 | 0.3611 | -0.0396 | 1127 | 1126 | 1 |
| 3CCW | 0.7652 | 0.5878 | 0.9969 | 0.5920 | 0.1124 | 549 | 541 | 8 |
| 3PBL | 0.6470 | 0.8780 | 0.9954 | 0.8748 | 0.0565 | 2228 | 2214 | 14 |
| 3BQD | 0.5618 | 0.7802 | 0.9949 | 0.7776 | 0.0212 | 998 | 992 | 6 |
| 3G0E | 0.8057 | 0.6887 | 0.9924 | 0.6899 | 0.1338 | 387 | 379 | 8 |
| 830C | 0.6763 | 0.7968 | 0.9902 | 0.7922 | 0.0906 | 1670 | 1644 | 26 |
| 2CNK | 0.7350 | 0.1928 | 0.9891 | 0.2495 | 0.1118 | 509 | 472 | 37 |
| 1XL2 | 0.8517 | 0.4639 | 0.9887 | 0.4910 | 0.1817 | 1607 | 1511 | 96 |
| 3EQH | 0.6921 | 0.2403 | 0.9867 | 0.2656 | 0.0704 | 320 | 308 | 12 |
| 2ZEC | 0.6770 | 0.3122 | 0.9857 | 0.3544 | 0.1373 | 237 | 221 | 16 |
| 2AM9 | 0.5055 | 0.8199 | 0.9835 | 0.8094 | 0.0103 | 1107 | 1088 | 19 |
| 1BCD | 0.4933 | 0.1675 | 0.9822 | 0.1753 | -0.0191 | 2002 | 1976 | 26 |
| 3D0E | 0.8424 | 0.6498 | 0.9809 | 0.6692 | 0.3015 | 260 | 237 | 23 |
| 2OI0 | 0.5505 | 0.5676 | 0.9808 | 0.5665 | 0.0250 | 1384 | 1353 | 31 |
| 1MV9 | 0.6055 | 0.8322 | 0.9806 | 0.8199 | 0.0465 | 311 | 304 | 7 |
| 3LPB | 0.4851 | 0.3690 | 0.9789 | 0.3760 | 0.0112 | 258 | 252 | 6 |
| 2QD9 | 0.6189 | 0.8174 | 0.9758 | 0.8036 | 0.0902 | 2291 | 2218 | 73 |
| 3HMM | 0.5769 | 0.7489 | 0.9670 | 0.7314 | -0.0420 | 242 | 235 | 7 |
| 2H7L | 0.5769 | 0.7489 | 0.9670 | 0.7314 | -0.0420 | 242 | 235 | 7 |
| 3L5D | 0.8266 | 0.9133 | 0.9665 | 0.8892 | 0.3445 | 641 | 600 | 41 |
| 3EML | 0.6269 | 0.4002 | 0.9665 | 0.4221 | 0.0847 | 3288 | 3096 | 192 |
| 2FSZ | 0.8597 | 0.9173 | 0.9661 | 0.8948 | 0.4686 | 1492 | 1366 | 126 |
| 2AYW | 0.7089 | 0.2182 | 0.9638 | 0.2946 | 0.1154 | 1093 | 976 | 117 |
| 3FRJ | 0.4572 | 0.1207 | 0.9633 | 0.1446 | -0.0093 | 899 | 870 | 29 |
| 2GTK | 0.3925 | 0.7785 | 0.9626 | 0.7564 | -0.0550 | 1334 | 1291 | 43 |
| 3LQ8 | 0.5867 | 0.6042 | 0.9621 | 0.6006 | 0.0583 | 353 | 336 | 17 |
| 1SJ0 | 0.8025 | 0.7057 | 0.9617 | 0.7078 | 0.2678 | 1451 | 1315 | 136 |
| 3BGS | 0.5118 | 0.8109 | 0.9602 | 0.7863 | 0.0055 | 248 | 238 | 10 |
| 3CJO | 0.6768 | 0.5109 | 0.9592 | 0.5377 | 0.1784 | 305 | 276 | 29 |
| 3CHP | 0.6478 | 0.8295 | 0.9567 | 0.8038 | 0.1265 | 367 | 346 | 21 |
| 2P2I | 0.6366 | 0.6983 | 0.9541 | 0.6840 | 0.0751 | 2462 | 2320 | 142 |
| 1UDT | 0.7414 | 0.7536 | 0.9531 | 0.7413 | 0.2310 | 1063 | 970 | 93 |
| 3D4Q | 0.7178 | 0.8202 | 0.9524 | 0.7971 | 0.2392 | 345 | 317 | 28 |
| 3KL6 | 0.5061 | 0.4785 | 0.9492 | 0.4817 | 0.0082 | 3340 | 3164 | 176 |
| 2ETR | 0.5402 | 0.6804 | 0.9490 | 0.6667 | 0.0766 | 234 | 219 | 15 |
| 1YPE | 0.6432 | 0.4269 | 0.9485 | 0.4636 | 0.1341 | 2541 | 2286 | 255 |
| 3HL5 | 0.5043 | 0.7300 | 0.9481 | 0.7103 | 0.0873 | 107 | 100 | 7 |
| 3BKL | 0.6001 | 0.6494 | 0.9479 | 0.6382 | 0.0621 | 868 | 813 | 55 |
| 3BIZ | 0.5069 | 0.9005 | 0.9476 | 0.8602 | 0.1302 | 236 | 221 | 15 |
| 2P54 | 0.6506 | 0.8819 | 0.9469 | 0.8441 | 0.1672 | 1174 | 1092 | 82 |
| 1SQT | 0.4882 | 0.1387 | 0.9455 | 0.2220 | 0.0640 | 419 | 375 | 44 |
| 2VT4 | 0.5656 | 0.4779 | 0.9429 | 0.5014 | 0.1192 | 726 | 657 | 69 |

| | | | | | | | | |
|---|---|---|---|---|---|---|---|---|
| 2HZI | 0.8138 | 0.6895 | 0.9400 | 0.7059 | 0.3660 | 493 | 409 | 84 |
| 3CQW | 0.6763 | 0.6037 | 0.9367 | 0.5991 | 0.0845 | 641 | 588 | 53 |
| 1B9V | 0.5898 | 0.5463 | 0.9333 | 0.5531 | 0.0962 | 226 | 205 | 21 |
| 2I78 | 0.5760 | 0.8865 | 0.9330 | 0.8364 | 0.0924 | 2194 | 2027 | 167 |
| 2RGP | 0.8190 | 0.7463 | 0.9322 | 0.7538 | 0.4423 | 2027 | 1620 | 407 |
| 1H00 | 0.6143 | 0.6033 | 0.9302 | 0.5992 | 0.0957 | 1462 | 1326 | 136 |
| 3G6Z | 0.6409 | 0.8668 | 0.9299 | 0.8221 | 0.2480 | 444 | 398 | 46 |
| 3F07 | 0.7875 | 0.8307 | 0.9298 | 0.8112 | 0.4865 | 392 | 319 | 73 |
| 2OF2 | 0.6871 | 0.5484 | 0.9265 | 0.5736 | 0.1923 | 1067 | 919 | 148 |
| 3E37 | 0.5940 | 0.5236 | 0.9249 | 0.5242 | 0.0298 | 1591 | 1459 | 132 |
| 1LI4 | 0.5538 | 0.6769 | 0.9167 | 0.6429 | -0.0683 | 70 | 65 | 5 |
| 2ICA | 0.6365 | 0.8025 | 0.9155 | 0.7507 | -0.0174 | 353 | 324 | 29 |
| 1C8K | 0.5284 | 0.2530 | 0.9130 | 0.2889 | -0.0201 | 90 | 83 | 7 |
| 3NXO | 0.4664 | 0.7588 | 0.8970 | 0.7023 | -0.0601 | 1253 | 1136 | 117 |
| 2ZNP | 0.5381 | 0.8696 | 0.8960 | 0.7917 | -0.0377 | 792 | 713 | 79 |
| 3MAX | 0.6439 | 0.8160 | 0.8915 | 0.7537 | 0.1293 | 475 | 413 | 62 |
| 2HV5 | 0.4378 | 0.2261 | 0.8896 | 0.2895 | 0.0062 | 684 | 606 | 78 |
| 1LRU | 0.2677 | 0.0925 | 0.8889 | 0.1215 | -0.1082 | 181 | 173 | 8 |
| 3BZ3 | 0.2466 | 0.8713 | 0.8889 | 0.7857 | -0.1196 | 112 | 101 | 11 |
| 2OJ9 | 0.5570 | 0.4316 | 0.8703 | 0.4732 | 0.0846 | 448 | 373 | 75 |
| 3EL8 | 0.6774 | 0.7549 | 0.8689 | 0.7071 | 0.2130 | 1560 | 1273 | 287 |
| 2ZDT | 0.4340 | 0.5495 | 0.8605 | 0.5156 | -0.1428 | 225 | 202 | 23 |
| 1E66 | 0.6548 | 0.5015 | 0.8551 | 0.5504 | 0.1826 | 2122 | 1635 | 487 |
| 1NJS | 0.5897 | 0.9808 | 0.8500 | 0.8438 | 0.3721 | 64 | 52 | 12 |
| 3LAN | 0.5831 | 0.6461 | 0.8490 | 0.6141 | 0.0877 | 1459 | 1201 | 258 |
| 1Q4X | 0.1732 | 0.5610 | 0.8415 | 0.5127 | -0.2101 | 275 | 246 | 29 |
| 3ODU | 0.7184 | 0.8372 | 0.8372 | 0.7544 | 0.3372 | 57 | 43 | 14 |
| 3NY8 | 0.4513 | 0.5622 | 0.8325 | 0.5373 | -0.0230 | 737 | 619 | 118 |
| 2NNQ | 0.8364 | 0.9362 | 0.8000 | 0.7778 | 0.3251 | 63 | 47 | 16 |
| 3M2W | 0.6844 | 0.6848 | 0.7826 | 0.6491 | 0.2384 | 265 | 184 | 81 |
| 1S3B | 0.4886 | 0.8377 | 0.7812 | 0.6906 | 0.0092 | 585 | 456 | 129 |
| 2I0E | 0.6469 | 0.6141 | 0.7793 | 0.6046 | 0.1795 | 521 | 368 | 153 |
| 2OJG | 0.5857 | 0.8148 | 0.7765 | 0.7069 | 0.2821 | 116 | 81 | 35 |
| 1SYN | 0.3875 | 0.2786 | 0.7724 | 0.3054 | -0.1812 | 465 | 402 | 63 |
| 1J4H | 0.5961 | 0.7818 | 0.7500 | 0.6609 | 0.1546 | 233 | 165 | 68 |
| 1UYG | 0.3655 | 0.6705 | 0.7375 | 0.5575 | -0.1548 | 113 | 88 | 25 |
| 2E1W | 0.5014 | 0.7404 | 0.7333 | 0.6233 | 0.0743 | 146 | 104 | 42 |
| 3C4F | 0.5703 | 0.6789 | 0.7115 | 0.5877 | 0.0609 | 473 | 327 | 146 |
| 3LN1 | 0.3907 | 0.2277 | 0.7099 | 0.3114 | -0.1247 | 2174 | 1730 | 444 |
| 1D3G | 0.4088 | 0.2247 | 0.7083 | 0.3746 | -0.0149 | 315 | 227 | 88 |
| 1KVO | 0.2119 | 0.0909 | 0.6957 | 0.1211 | -0.3277 | 190 | 176 | 14 |
| 1W7X | 0.2159 | 0.1508 | 0.6866 | 0.1860 | -0.3103 | 344 | 305 | 39 |
| 3KGC | 0.3955 | 0.5123 | 0.6631 | 0.4702 | -0.1092 | 689 | 488 | 201 |
| 2V3F | 0.3983 | 0.4000 | 0.6471 | 0.4512 | -0.0424 | 82 | 55 | 27 |

| | | | | | | | | |
|---|---|---|---|---|---|---|---|---|
| 2OYU | 0.5191 | 0.0976 | 0.6023 | 0.6745 | 0.1350 | 1613 | 543 | 1070 |
| 2AZR | 0.4730 | 0.0458 | 0.4815 | 0.3478 | -0.0906 | 437 | 284 | 153 |
| 3NF7 | 0.5717 | 0.6973 | 0.4388 | 0.5121 | 0.0841 | 453 | 185 | 268 |
| 1L2S | 0.6311 | 0.5510 | 0.4355 | 0.5714 | 0.1299 | 133 | 49 | 84 |
| 3NXU | 0.3981 | 0.3135 | 0.4241 | 0.4088 | -0.1733 | 570 | 303 | 267 |
| 1QW6 | 0.4563 | 0.0155 | 0.3846 | 0.1772 | -0.2046 | 395 | 322 | 73 |
| 1R9O | 0.3632 | 0.1172 | 0.3617 | 0.5078 | -0.0749 | 321 | 145 | 176 |
| 1VSO | 0.3137 | 0.4706 | 0.2712 | 0.3423 | -0.2617 | 371 | 136 | 235 |
| 2B8T | 0.4948 | 0.0000 | nan | 0.5397 | 0.0000 | 126 | 58 | 68 |
| 3F9M | 0.5357 | 0.0000 | nan | 0.0886 | 0.0000 | 158 | 144 | 14 |

**Table S4. The performance of DeepBindGCN_RG(pocket cutoff 0.8nm) on the DUD.E dataset.**

| PDBID | Rmse | Mse | Pearson | Spearman | CI | data_size |
|---|---|---|---|---|---|---|
| 3BIZ | 0.6866 | 0.4714 | 0.1794 | 0.1800 | 0.5570 | 221 |
| 2AZR | 0.7134 | 0.5089 | 0.2293 | 0.2654 | 0.5903 | 284 |
| 1UYG | 0.7880 | 0.6209 | 0.3155 | 0.2981 | 0.6089 | 88 |
| 3M2W | 0.7958 | 0.6334 | 0.3754 | 0.3063 | 0.6073 | 184 |
| 3EQH | 0.8114 | 0.6584 | 0.3547 | 0.3277 | 0.6159 | 308 |
| 2ETR | 0.8119 | 0.6592 | 0.2780 | 0.2687 | 0.5961 | 219 |
| 3F9M | 0.8177 | 0.6686 | 0.1705 | 0.1740 | 0.5611 | 144 |
| 1KVO | 0.8184 | 0.6697 | 0.1789 | 0.1481 | 0.5510 | 176 |
| 1SQT | 0.8194 | 0.6715 | 0.2473 | 0.2282 | 0.5777 | 375 |
| 3D0E | 0.8439 | 0.7122 | 0.2704 | 0.2272 | 0.5797 | 237 |
| 3L5D | 0.8480 | 0.7191 | 0.3180 | 0.3432 | 0.6187 | 600 |
| 1LRU | 0.8956 | 0.8021 | 0.2213 | 0.2362 | 0.5805 | 173 |
| 3NF7 | 0.9010 | 0.8119 | 0.1790 | 0.1021 | 0.5353 | 185 |
| 3HMM | 0.9035 | 0.8163 | 0.0380 | 0.0055 | 0.5010 | 235 |
| 2ICA | 0.9056 | 0.8201 | 0.3269 | 0.3630 | 0.6210 | 324 |
| 2HZI | 0.9088 | 0.8258 | 0.5412 | 0.5701 | 0.6958 | 409 |
| 3KGC | 0.9121 | 0.8319 | -0.0222 | 0.0049 | 0.5013 | 488 |
| 2HV5 | 0.9258 | 0.8572 | 0.0512 | 0.0530 | 0.5178 | 606 |
| 3EL8 | 0.9303 | 0.8654 | 0.2629 | 0.2570 | 0.5875 | 1271 |
| 2OJG | 0.9386 | 0.8810 | 0.5505 | 0.5713 | 0.7045 | 81 |
| 1D3G | 0.9397 | 0.8831 | 0.0503 | 0.0742 | 0.5269 | 227 |
| 1BCD | 0.9496 | 0.9017 | 0.3138 | 0.2846 | 0.5974 | 1976 |
| 2V3F | 0.9621 | 0.9256 | 0.3420 | 0.2885 | 0.5987 | 55 |
| 3CCW | 0.9665 | 0.9341 | 0.2556 | 0.2955 | 0.6004 | 541 |
| 2QD9 | 0.9730 | 0.9468 | 0.3492 | 0.3509 | 0.6196 | 2218 |
| 3KRJ | 0.9770 | 0.9545 | 0.2654 | 0.2395 | 0.5826 | 389 |
| 3CQW | 0.9779 | 0.9562 | 0.2804 | 0.2742 | 0.5933 | 588 |
| 2ZNP | 0.9779 | 0.9564 | 0.1656 | 0.1517 | 0.5510 | 713 |
| 2OF2 | 0.9816 | 0.9635 | 0.2678 | 0.2355 | 0.5797 | 919 |

| | | | | | | |
|---|---|---|---|---|---|---|
| 830C | 0.9833 | 0.9668 | 0.2000 | 0.1883 | 0.5641 | 1644 |
| 3LAN | 0.9854 | 0.9709 | 0.1809 | 0.1732 | 0.5596 | 1201 |
| 2OJ9 | 0.9918 | 0.9836 | 0.4426 | 0.4041 | 0.6388 | 373 |
| 3MAX | 0.9936 | 0.9873 | 0.0286 | 0.0379 | 0.5130 | 413 |
| 1J4H | 0.9965 | 0.9930 | -0.1850 | -0.1821 | 0.4383 | 165 |
| 3G0E | 0.9967 | 0.9935 | 0.0037 | -0.0001 | 0.4966 | 379 |
| 1UDT | 0.9988 | 0.9976 | 0.4255 | 0.4115 | 0.6419 | 970 |
| 3FRJ | 1.0053 | 1.0107 | 0.3219 | 0.3558 | 0.6219 | 870 |
| 3LN1 | 1.0114 | 1.0229 | 0.1226 | 0.1406 | 0.5467 | 1724 |
| 2OYU | 1.0176 | 1.0356 | 0.0176 | -0.0037 | 0.4991 | 542 |
| 1MV9 | 1.0219 | 1.0443 | -0.0451 | -0.0704 | 0.4770 | 302 |
| 2I0E | 1.0248 | 1.0503 | 0.1046 | 0.0637 | 0.5211 | 368 |
| 3G6Z | 1.0335 | 1.0681 | 0.3708 | 0.3808 | 0.6315 | 398 |
| 2P54 | 1.0358 | 1.0729 | -0.0942 | -0.0807 | 0.4732 | 1092 |
| 3C4F | 1.0446 | 1.0912 | 0.2187 | 0.2043 | 0.5695 | 327 |
| 2OI0 | 1.0482 | 1.0987 | 0.2427 | 0.2715 | 0.5921 | 1353 |
| 3BZ3 | 1.0552 | 1.1134 | 0.4161 | 0.2853 | 0.5996 | 101 |
| 2FSZ | 1.0561 | 1.1154 | 0.1761 | 0.1903 | 0.5635 | 1366 |
| 3L3M | 1.0662 | 1.1368 | 0.2866 | 0.2893 | 0.5993 | 1045 |
| 2I78 | 1.0709 | 1.1468 | 0.2367 | 0.2588 | 0.5878 | 2027 |
| 2NNQ | 1.0723 | 1.1498 | -0.0430 | 0.0930 | 0.5363 | 47 |
| 1W7X | 1.0802 | 1.1669 | -0.1663 | -0.1278 | 0.4586 | 305 |
| 1H00 | 1.0811 | 1.1687 | 0.1471 | 0.1481 | 0.5504 | 1326 |
| 3CJO | 1.1029 | 1.2165 | 0.0636 | 0.0040 | 0.5023 | 276 |
| 2GTK | 1.1122 | 1.2370 | 0.1875 | 0.1640 | 0.5554 | 1291 |
| 3E37 | 1.1124 | 1.2374 | 0.3073 | 0.2869 | 0.5971 | 1458 |
| 3LQ8 | 1.1194 | 1.2531 | 0.2573 | 0.2449 | 0.5826 | 336 |
| 2AA2 | 1.1231 | 1.2614 | 0.1248 | 0.1253 | 0.5415 | 212 |
| 3BQD | 1.1239 | 1.2632 | 0.2318 | 0.2555 | 0.5849 | 992 |
| 2P2I | 1.1261 | 1.2682 | 0.1844 | 0.1634 | 0.5554 | 2320 |
| 3KL6 | 1.1401 | 1.2998 | 0.3700 | 0.3639 | 0.6240 | 3164 |
| 3D4Q | 1.1457 | 1.3127 | 0.3178 | 0.3248 | 0.6096 | 317 |
| 3NXU | 1.1501 | 1.3226 | 0.1605 | 0.0808 | 0.5280 | 301 |
| 1LI4 | 1.1507 | 1.3240 | 0.1877 | 0.1849 | 0.5623 | 65 |
| 3KBA | 1.1584 | 1.3418 | 0.2295 | 0.1383 | 0.5450 | 1126 |
| 3BKL | 1.1908 | 1.4180 | 0.1231 | 0.1792 | 0.5606 | 813 |
| 3CHP | 1.2059 | 1.4543 | 0.2302 | 0.2447 | 0.5830 | 346 |
| 3F07 | 1.2073 | 1.4576 | 0.0710 | 0.0406 | 0.5144 | 319 |
| 2ZEC | 1.2095 | 1.4629 | 0.1899 | 0.2193 | 0.5701 | 221 |
| 2OWB | 1.2178 | 1.4830 | -0.0745 | 0.0285 | 0.5094 | 227 |
| 1SYN | 1.2312 | 1.5158 | 0.1394 | 0.1719 | 0.5579 | 402 |
| 2ZDT | 1.2347 | 1.5244 | 0.0851 | 0.0698 | 0.5228 | 202 |
| 3PBL | 1.2370 | 1.5302 | 0.2382 | 0.2288 | 0.5772 | 2212 |
| 3NY8 | 1.2412 | 1.5405 | 0.0940 | 0.0653 | 0.5224 | 612 |

| | | | | | | |
|---|---|---|---|---|---|---|
| 2CNK | 1.2492 | 1.5606 | 0.1769 | 0.2146 | 0.5712 | 472 |
| 1VSO | 1.2498 | 1.5619 | -0.0160 | -0.0015 | 0.5018 | 136 |
| 1R9O | 1.2565 | 1.5787 | 0.0482 | -0.0124 | 0.4960 | 143 |
| 2AYW | 1.2632 | 1.5957 | 0.2006 | 0.2593 | 0.5870 | 975 |
| 1E66 | 1.2722 | 1.6184 | 0.0327 | 0.0638 | 0.5212 | 1635 |
| 3LPB | 1.2798 | 1.6379 | 0.2051 | 0.2103 | 0.5699 | 252 |
| 1XL2 | 1.2904 | 1.6652 | 0.2034 | 0.2053 | 0.5695 | 1510 |
| 2RGP | 1.3472 | 1.8149 | -0.0431 | -0.0291 | 0.4903 | 1620 |
| 1C8K | 1.3568 | 1.8408 | -0.2455 | -0.1909 | 0.4346 | 83 |
| 2AM9 | 1.3654 | 1.8643 | 0.0073 | 0.0170 | 0.5063 | 1086 |
| 1SJ0 | 1.3782 | 1.8994 | 0.2670 | 0.2893 | 0.5966 | 1315 |
| 1ZW5 | 1.3800 | 1.9045 | -0.0774 | -0.1686 | 0.4433 | 85 |
| 1L2S | 1.4351 | 2.0594 | 0.3744 | 0.2842 | 0.6015 | 49 |
| 1Q4X | 1.5252 | 2.3263 | 0.1688 | 0.1581 | 0.5511 | 246 |
| 3EML | 1.5608 | 2.4360 | 0.2073 | 0.1968 | 0.5664 | 3094 |
| 2E1W | 1.5950 | 2.5441 | 0.0564 | 0.3352 | 0.6280 | 104 |
| 3BWM | 1.6102 | 2.5927 | 0.5564 | 0.5051 | 0.6847 | 41 |
| 2VT4 | 1.6758 | 2.8084 | -0.0898 | -0.1105 | 0.4632 | 656 |
| 3NXO | 1.7128 | 2.9336 | 0.1981 | 0.2121 | 0.5701 | 1136 |
| 3BGS | 1.7390 | 3.0243 | 0.2201 | 0.2084 | 0.5703 | 238 |
| 1B9V | 1.7737 | 3.1460 | -0.0314 | -0.0547 | 0.4799 | 205 |
| 3ODU | 1.8775 | 3.5251 | -0.5341 | -0.5619 | 0.3000 | 43 |
| 1YPE | 1.9325 | 3.7347 | 0.1137 | 0.0743 | 0.5261 | 2286 |
| 2H7L | 1.9446 | 3.7814 | -0.0870 | -0.1644 | 0.4295 | 44 |
| 2B8T | 2.0177 | 4.0713 | 0.4273 | 0.3408 | 0.6174 | 58 |
| 1QW6 | 2.1217 | 4.5014 | 0.0362 | 0.0522 | 0.5173 | 322 |
| 1NJS | 2.2257 | 4.9538 | -0.0284 | -0.0702 | 0.4784 | 52 |
| 3HL5 | 2.3382 | 5.4670 | 0.1609 | 0.2207 | 0.5754 | 100 |
| 1S3B | 2.6465 | 7.0038 | -0.1133 | -0.0348 | 0.4883 | 455 |
| Average | 1.1893 | 1.5440 | 0.1655 | 0.1626 | 0.5557 | 668.8725 |

**Table S5**. The top predicted candidates from DeepBindGCN_BC and DeepBindGCN_RG for the PD-L1 dimer.

| **Compound ID** | **DeepBindGCN_BC** | **DeepBindGCN_RG** | **Schrödinger score** |
|---|---:|---:|---:|
| M769-0298 | 1.0000 | 8.6240 | -7.0316 |
| 8510-0254 | 1.0000 | 8.6550 | -8.5080 |
| 7840-3923 | 1.0000 | 8.6628 | -8.9960 |
| K305-0238 | 1.0000 | 8.7054 | None |
| M769-0279 | 1.0000 | 8.7637 | -8.0958 |
| E955-1714 | 1.0000 | 8.8704 | None |
| K286-3615 | 1.0000 | 8.6922 | -6.2060 |
| P181-0863 | 1.0000 | 8.6661 | -7.0520 |

| | | | |
|---|---|---|---|
| E955-1146 | 1.0000 | 8.6790 | None |
| 4376-0091 | 1.0000 | 8.6287 | -9.5048 |
| P491-4168 | 1.0000 | 8.7444 | -4.4769 |
| P491-4131 | 1.0000 | 8.7610 | -7.1448 |
| P491-4650 | 1.0000 | 8.6005 | -6.7671 |
| 4296-0364 | 1.0000 | 8.6700 | None |
| L880-0302 | 1.0000 | 8.6232 | -5.7479 |
| C163-0031 | 1.0000 | 9.0551 | -7.3762 |
| F019-0136 | 1.0000 | 8.9002 | -5.3502 |
| E955-1004 | 1.0000 | 9.1059 | None |
| K305-0239 | 1.0000 | 8.6878 | None |
| E955-1572 | 1.0000 | 8.7449 | None |
| V010-3729 | 1.0000 | 8.6142 | -9.5620 |
| C301-3800 | 1.0000 | 8.6228 | None |
| G833-0486 | 0.9999 | 8.6179 | -8.4276 |
| 7418-0675 | 0.9999 | 8.6696 | -8.5439 |
| M769-0266 | 0.9999 | 8.9623 | -8.9308 |
| 0957-0218 | 0.9999 | 8.6395 | -9.1960 |
| M769-1241 | 0.9999 | 8.7414 | -9.2589 |
| K286-3705 | 0.9999 | 8.6012 | -5.9413 |
| L880-0055 | 0.9999 | 8.6495 | -7.0515 |
| E955-0152 | 0.9999 | 8.7428 | None |
| 1890-0396 | 0.9999 | 8.6412 | -7.6736 |
| E955-0720 | 0.9999 | 8.6566 | None |
| G281-2300 | 0.9999 | 8.6932 | -2.7696 |
| K305-0345 | 0.9999 | 8.6874 | None |
| F506-0470 | 0.9999 | 8.6808 | None |
| P491-4166 | 0.9999 | 8.6334 | -6.9812 |
| 6049-2376 | 0.9999 | 8.9938 | -6.9023 |
| M769-0273 | 0.9999 | 8.6812 | -8.5714 |
| L880-0048 | 0.9999 | 8.6338 | -5.7588 |
| M769-0271 | 0.9999 | 8.8637 | -6.3845 |
| 3209-0888 | 0.9998 | 8.6119 | -6.8953 |
| M769-0296 | 0.9998 | 8.6302 | -7.7200 |
| M769-0268 | 0.9998 | 8.8134 | -4.7701 |
| M769-0256 | 0.9998 | 8.7986 | -8.5208 |
| F940-0602 | 0.9997 | 8.6125 | -8.9528 |
| D725-0206 | 0.9997 | 8.6493 | -8.5324 |
| G822-0592 | 0.9997 | 8.6111 | -5.7201 |
| C163-0098 | 0.9996 | 8.7322 | -8.3344 |
| 6049-1532 | 0.9996 | 8.7974 | -6.2506 |
| K305-0045 | 0.9996 | 8.6580 | None |
| C200-0820 | 0.9996 | 8.7744 | None |
| L558-0642 | 0.9995 | 8.6285 | None |

| | | | |
|---|---|---|---|
| E859-0698 | 0.9995 | 8.6636 | -9.5635 |
| K286-3702 | 0.9995 | 8.6177 | None |
| E955-0436 | 0.9995 | 8.7842 | None |
| K305-0044 | 0.9994 | 8.7119 | None |
| 3852-0327 | 0.9994 | 8.6303 | -8.1867 |
| 7418-1976 | 0.9994 | 8.6795 | -9.0079 |
| L558-0647 | 0.9994 | 8.7484 | None |
| M460-2243 | 0.9993 | 8.6750 | -7.3363 |
| F834-0590 | 0.9993 | 8.6778 | -8.7997 |
| V001-2516 | 0.9992 | 8.6688 | -9.6745 |
| L880-0269 | 0.9992 | 8.6402 | -8.2118 |
| L880-0296 | 0.9992 | 8.7370 | -4.7731 |
| Y043-4611 | 0.9992 | 8.6051 | -7.6287 |
| V006-3291 | 0.9992 | 8.7025 | -9.4725 |
| M460-2244 | 0.9991 | 8.9798 | -7.8858 |
| 3209-0884 | 0.9990 | 8.6122 | None |
| L855-0058 | 0.9989 | 8.6996 | -4.5421 |
| M769-1425 | 0.9989 | 8.7582 | -7.3411 |
| C200-0812 | 0.9987 | 8.9092 | -6.5423 |
| G281-2304 | 0.9986 | 8.6122 | -1.3766 |
| 8010-2605 | 0.9985 | 8.6211 | None |
| G856-8325 | 0.9984 | 8.7683 | -9.6361 |
| C163-0025 | 0.9981 | 8.8068 | -8.0547 |
| V030-9758 | 0.9981 | 8.9498 | -8.9287 |
| V006-3679 | 0.9979 | 8.7418 | -9.2080 |
| M460-2265 | 0.9978 | 8.6296 | -8.4071 |
| G856-8308 | 0.9977 | 8.6416 | -9.0176 |
| L880-0285 | 0.9976 | 8.7407 | -5.1432 |
| P704-0166 | 0.9974 | 8.6151 | -7.4346 |
| L539-0325 | 0.9973 | 8.6117 | -7.8659 |
| L977-1354 | 0.9973 | 8.6063 | -2.7656 |
| 8007-8597 | 0.9972 | 8.6505 | -7.5859 |
| 2265-3136 | 0.9972 | 8.6034 | None |
| M520-0742 | 0.9970 | 8.6904 | -8.8531 |
| V008-6168 | 0.9968 | 8.8348 | -8.5468 |
| M520-0662 | 0.9967 | 8.6738 | -8.7562 |
| 8015-4975 | 0.9964 | 8.7747 | -5.5618 |
| C226-4308 | 0.9960 | 8.8256 | -7.5766 |
| E955-0805 | 0.9949 | 8.6627 | None |
| L880-0286 | 0.9948 | 8.6384 | -5.7523 |
| F431-0440 | 0.9943 | 8.6648 | -7.3802 |
| Y020-7930 | 0.9941 | 8.6259 | -5.6173 |
| M460-3727 | 0.9939 | 8.6979 | -8.5876 |
| P392-2143 | 0.9933 | 8.7914 | -9.0883 |

| | | | |
|---|---|---|---|
| L858-0205 | 0.9932 | 8.6213 | -3.2103 |
| M460-2223 | 0.9926 | 8.6092 | -6.9175 |
| M460-2191 | 0.9923 | 8.6177 | -6.6068 |
| J093-0740 | 0.9922 | 8.6521 | -8.4309 |
| M769-1402 | 0.9922 | 8.6924 | -6.9686 |
| SB33-0022 | 0.9917 | 8.6459 | -6.0260 |
| S556-0709 | 0.9917 | 8.7039 | -7.0693 |
| M520-0729 | 0.9914 | 8.8291 | -9.1131 |
| C163-0043 | 0.9909 | 8.6221 | -8.2081 |
| K305-0042 | 0.9908 | 8.8660 | -6.9674 |
| V008-7569 | 0.9902 | 8.7126 | -5.2661 |
| E955-1055 | 0.9901 | 8.6774 | None |
| D725-0060 | 0.9900 | 8.6192 | -6.8281 |

**Table S6. The performance of DeepBindGCN_RG_x with different training epoch on the PDBbind v.2016 core set (CASF-2016).**

| Epoch | Rmse | Mse | Pearson | Spearman | CI |
|---|---|---|---|---|---|
| 100 | 1.4491 | 2.0998 | 0.7593 | 0.7574 | 0.7826 |
| 200 | 1.4500 | 2.1024 | 0.7508 | 0.7577 | 0.7857 |
| 300 | 1.4268 | 2.0357 | 0.7605 | 0.7615 | 0.7866 |
| 400 | 1.3874 | 1.9250 | 0.7699 | 0.7692 | 0.7902 |
| 500 | 1.4299 | 2.0447 | 0.7534 | 0.7542 | 0.7815 |
| 600 | 1.4279 | 2.0390 | 0.7539 | 0.7525 | 0.7807 |
| 700 | 1.4240 | 2.0279 | 0.7588 | 0.7570 | 0.7848 |
| 800 | 1.3969 | 1.9514 | 0.7696 | 0.7593 | 0.7859 |
| 900 | 1.4007 | 1.9620 | 0.7655 | 0.7618 | 0.7849 |
| 1000 | 1.3973 | 1.9524 | 0.7659 | 0.7530 | 0.7810 |
| 1100 | 1.3867 | 1.9230 | 0.7703 | 0.7642 | 0.7855 |
| 1200 | 1.3843 | 1.9162 | 0.7719 | 0.7672 | 0.7891 |
| 1300 | 1.3882 | 1.9270 | 0.7696 | 0.7573 | 0.7847 |
| 1400 | 1.3981 | 1.9547 | 0.7656 | 0.7533 | 0.7833 |
| 1500 | 1.4094 | 1.9864 | 0.7610 | 0.7440 | 0.7768 |

| | | | | | |
|---|---|---|---|---|---|
| 1600 | 1.4304 | 2.0461 | 0.7533 | 0.7405 | 0.7757 |
| 1700 | 1.4292 | 2.0426 | 0.7538 | 0.7407 | 0.7772 |
| 1800 | 1.4322 | 2.0511 | 0.7520 | 0.7369 | 0.7750 |
| 1900 | 1.3985 | 1.9558 | 0.7694 | 0.7538 | 0.7814 |
| 2000 | 1.4190 | 2.0136 | 0.7584 | 0.7431 | 0.7773 |

**Table S7. The performance of DeepBindGCN_RG_x with different training epoch on the PDBbind v.2013 core set.**

| Epoch | Rmse | Mse | Pearson | Spearman | CI |
|---|---|---|---|---|---|
| 100 | 1.6113 | 2.5963 | 0.7078 | 0.7019 | 0.7563 |
| 200 | 1.5519 | 2.4083 | 0.7340 | 0.7259 | 0.7677 |
| 300 | 1.5121 | 2.2864 | 0.7415 | 0.7216 | 0.7686 |
| 400 | 1.4921 | 2.2263 | 0.7493 | 0.7372 | 0.7722 |
| 500 | 1.5385 | 2.3668 | 0.7295 | 0.7166 | 0.7640 |
| 600 | 1.5103 | 2.2811 | 0.7413 | 0.7306 | 0.7730 |
| 700 | 1.5064 | 2.2693 | 0.7460 | 0.7366 | 0.7756 |
| 800 | 1.4931 | 2.2294 | 0.7506 | 0.7289 | 0.7726 |
| 900 | 1.4828 | 2.1986 | 0.7514 | 0.7321 | 0.7741 |
| 1000 | 1.4823 | 2.1971 | 0.7524 | 0.7339 | 0.7734 |
| 1100 | 1.4750 | 2.1756 | 0.7548 | 0.7417 | 0.7758 |
| 1200 | 1.4864 | 2.2094 | 0.7503 | 0.7358 | 0.7740 |
| 1300 | 1.5318 | 2.3463 | 0.7314 | 0.7153 | 0.7667 |
| 1400 | 1.4775 | 2.1830 | 0.7539 | 0.7390 | 0.7779 |
| 1500 | 1.5091 | 2.2775 | 0.7414 | 0.7222 | 0.7669 |
| 1600 | 1.5226 | 2.3183 | 0.7367 | 0.7219 | 0.7665 |
| 1700 | 1.5351 | 2.3564 | 0.7307 | 0.7170 | 0.7659 |
| 1800 | 1.5011 | 2.2534 | 0.7450 | 0.7224 | 0.7687 |
| 1900 | 1.4844 | 2.2035 | 0.7540 | 0.7375 | 0.7743 |
| 2000 | 1.4986 | 2.2458 | 0.7467 | 0.7272 | 0.7725 |

**Table S8. We listed performance of DeepBindGCN_BC_onehot representation on test set during the training with epoch interval 100.**

| Epoch | AUC | TPR | Precision | Accuracy |
|---|---|---|---|---|
| 100 | 0.8420 | 0.6208 | 0.5950 | 0.7996 |
| 200 | 0.8509 | 0.6088 | 0.6185 | 0.8083 |
| 300 | 0.8586 | 0.6238 | 0.6204 | 0.8105 |
| 400 | 0.8615 | 0.6017 | 0.6452 | 0.8177 |
| 500 | 0.8607 | 0.6075 | 0.6353 | 0.8147 |
| 600 | 0.8662 | 0.6383 | 0.6289 | 0.8154 |
| 700 | 0.8684 | 0.6021 | 0.6653 | 0.8248 |
| 800 | 0.8663 | 0.6263 | 0.6555 | 0.8243 |
| 900 | 0.8672 | 0.6575 | 0.6345 | 0.8197 |
| 1000 | 0.8722 | 0.6208 | 0.6709 | 0.8291 |
| 1100 | 0.8722 | 0.6363 | 0.6628 | 0.8281 |
| 1200 | 0.8740 | 0.6342 | 0.6708 | 0.8307 |
| 1300 | 0.8748 | 0.6254 | 0.6692 | 0.8291 |
| 1400 | 0.8758 | 0.6388 | 0.6721 | 0.8318 |
| 1500 | 0.8704 | 0.6038 | 0.6806 | 0.8301 |
| 1600 | 0.8697 | 0.6442 | 0.6526 | 0.8253 |
| 1700 | 0.8719 | 0.6400 | 0.6615 | 0.8281 |
| 1800 | 0.8744 | 0.6421 | 0.6668 | 0.8303 |
| 1900 | 0.8762 | 0.6746 | 0.6489 | 0.8274 |
| 2000 | 0.8667 | 0.6471 | 0.6642 | 0.8300 |

**Table S9. We listed performance of DeepBindGCN_RG_onehot on test set during the training with epoch interval 100.**

| Epoch | Rmse | Mse | Pearson | Spearman | CI |
|---|---|---|---|---|---|
| 100 | 1.4180 | 2.0108 | 0.6785 | 0.6717 | 0.7438 |
| 200 | 1.5076 | 2.2728 | 0.6620 | 0.6531 | 0.7365 |
| 300 | 1.4459 | 2.0907 | 0.6772 | 0.6663 | 0.7416 |
| 400 | 1.4240 | 2.0278 | 0.6813 | 0.6703 | 0.7433 |
| 500 | 1.3913 | 1.9357 | 0.6904 | 0.6798 | 0.7475 |
| 600 | 1.3880 | 1.9266 | 0.6920 | 0.6789 | 0.7474 |
| 700 | 1.3441 | 1.8067 | 0.7066 | 0.6954 | 0.7548 |
| 800 | 1.3422 | 1.8015 | 0.7072 | 0.6941 | 0.7541 |
| 900 | 1.3364 | 1.7860 | 0.7147 | 0.7052 | 0.7591 |
| 1000 | 1.3249 | 1.7553 | 0.7146 | 0.7025 | 0.7584 |

| | | | | | |
|---|---|---|---|---|---|
| 1100 | 1.3130 | 1.7241 | 0.7238 | 0.7123 | 0.7625 |
| 1200 | 1.3006 | 1.6915 | 0.7249 | 0.7125 | 0.7635 |
| 1300 | 1.3008 | 1.6920 | 0.7246 | 0.7129 | 0.7634 |
| 1400 | 1.3153 | 1.7299 | 0.7179 | 0.7031 | 0.7589 |
| 1500 | 1.3075 | 1.7095 | 0.7226 | 0.7072 | 0.7613 |
| 1600 | 1.3154 | 1.7303 | 0.7155 | 0.6988 | 0.7574 |
| 1700 | 1.3076 | 1.7099 | 0.7196 | 0.7024 | 0.7589 |
| 1800 | 1.3051 | 1.7034 | 0.7223 | 0.7070 | 0.7610 |
| 1900 | 1.2987 | 1.6865 | 0.7247 | 0.7088 | 0.7613 |
| 2000 | 1.3060 | 1.7058 | 0.7231 | 0.7062 | 0.7607 |
